# Selfish chromosomal drive shapes recent centromeric histone evolution in monkeyflowers

**DOI:** 10.1101/2020.09.11.293597

**Authors:** Findley R. Finseth, Thom C. Nelson, Lila Fishman

## Abstract

Under the selfish centromere model, costs associated with female meiotic drive by centromeres select on interacting kinetochore proteins to restore Mendelian inheritance. We directly test this model in yellow monkeyflowers (*Mimulus guttatus*), which are polymorphic for a costly driving centromere (*D*). We show that the *D* haplotype is structurally and genetically distinct and swept to a high stable frequency within the past 1500 years. Quantitative genetic analyses reveal that variation in the strength of drive primarily depends on the identity of the non-*D* centromere, but also identified an unlinked modifier coincident with kinetochore protein Centromere-specific Histone 3 A (CenH3A). CenH3A has also experienced a recent (<1000 years) selective sweep in our focal population, consistent with ongoing interactions with *D* shaping its evolution. Together, our results demonstrate an active co-evolutionary arms race between the DNA and protein components of the meiotic machinery, with important consequences for individual fitness and molecular divergence.

## INTRODUCTION

Centromeres, which mediate the conserved and essential processes of chromosomal segregation during eukaryotic mitosis and meiosis, are paradoxically diverse. Centromeric DNA arrays are highly variable in sequence, size, and position, and the protein that epigenetically marks the site of kinetochore assembly, Centromere-specific Histone 3 (CenH3; known as CENP-A in humans), commonly evolves under diversifying selection (Malik and Henikoff 2001; Henikoff et al. 2001; Finseth et al. 2015). The selfish centromere hypothesis (Henikoff et al. 2001; Malik and Henikoff 2002) resolves this paradox by arguing: a) asymmetric female meiosis creates an arena for selection among homologous centromeres for inclusion in the single egg cell, b) female meiotic drive is costly to individuals and c) costs of drive promote suppressive coevolution by CenH3 and other key kinetochore proteins. This model of genetic conflict between the DNA and protein components of centromeres has profound implications for the maintenance of individual fitness variation, the divergence of species, and the evolution of genomes and cellular processes (McLaughlin and Malik 2017; Lampson and Black 2017; reviewed in Kursel and Malik 2018). Furthermore, understanding centromere function and evolution directly impact human endeavors from cancer therapies (Zhang et al. 2016) to crop improvement (Ravi and Chan 2010). However, despite recent advances in understanding the molecular biology (Chmátal et al. 2014; Akera et al. 2017) and evolutionary dynamics (Fishman and Kelly 2015) of centromeric drive, evidence for the posited evolutionary arms race between centromere DNA and kinetochore proteins remains largely circumstantial. Here, we directly test the key final step of the centromere drive hypothesis in a flowering plant with an active (and costly) driving centromere.

In the yellow monkeyflower, *Mimulus guttatus*, the *D* allele on Linkage Group 11 (LG11) drives through female meiosis against both conspecific (*M. guttatus D^−^* allele; *D:D*^−^ female transmission ratio = 58:42) and heterospecific (M*. nasutus d* allele; *D:d* ratio > 98:2) alternative alleles in hybrids (Fishman and Willis 2005; Fishman and Saunders 2008). *D* is genetically and cytogenetically associated with dramatically expanded arrays of the *M. guttatus* centromere-specific DNA repeat Cent728 (Fishman and Saunders 2008; Melters et al. 2013). Along with near-perfect transmission in heterospecific F_1_ hybrids, which is only possible via centromeric drive in Meiosis I (Malik 2005), this association strongly suggests that *D* acts as the functional centromere of LG11. Homozygous costs to both male (pollen viability) and female fertility (seeds per fruit) prevent fixation of *D*, maintaining it at 35-45% in our focal annual Iron Mountain (IM) *M. guttatus* population (Oregon Cascades, USA) (Fishman and Saunders 2008; Fishman and Kelly 2015). Recent genome-wide association mapping of flowering traits in the field found little or no effect of *D* on other fitness components (Troth et al. 2018), confirming that its evolutionary dynamics primarily reflect a balance between selfish female meiotic drive and fertility costs. Because a costly driver at a polymorphic equilibrium generates selection for unlinked suppressor loci (Crow 1991), this population provides the ideal opportunity to assess the consequences of centromeric drive for selection on linked and unlinked genes.

## RESULTS

Comparative linkage mapping demonstrates local suppression of recombination in F_1_ hybrids of the IM62 *M. guttatus* reference line (*D*) with *D*^−^ and *d* lines (Flagel et al. 2019), suggesting that the *Cent728* expansions associated with *D* are embedded in a chromosomal rearrangement (likely an inversion) that reduces chromosomal pairing or crossing over between alternative haplotypes. Because the *M. guttatus* reference genome sequence was assembled into chromosome-scale scaffolds using a locally non-informative *D* × *D*^−^ linkage map, we generated a corrected LG11 genome order based on a collinear *D*^−^ × *D*^−^ map (Table S1) (Flagel et al. 2019). Using this collinear (but likely inverted relative to *D* chromosomes) order, IM inbred lines exhibit a contiguous block of elevated linkage disequilibrium (LD) across the region of LG11 corresponding to the driving *D* haplotype (Meiotic Drive Locus 11 or MDL11: Fig. 1; Fig. S1A; Table S2). This >12 Mb block almost certainly underestimates the true physical extent of MDL11; although this region contains extensive arrays of Cent728 repeats (Fig. 1), repetitive centromeric and peri-centromeric DNA are likely under-represented in the assembled and mapped genome scaffolds.

**Fig. 1.**
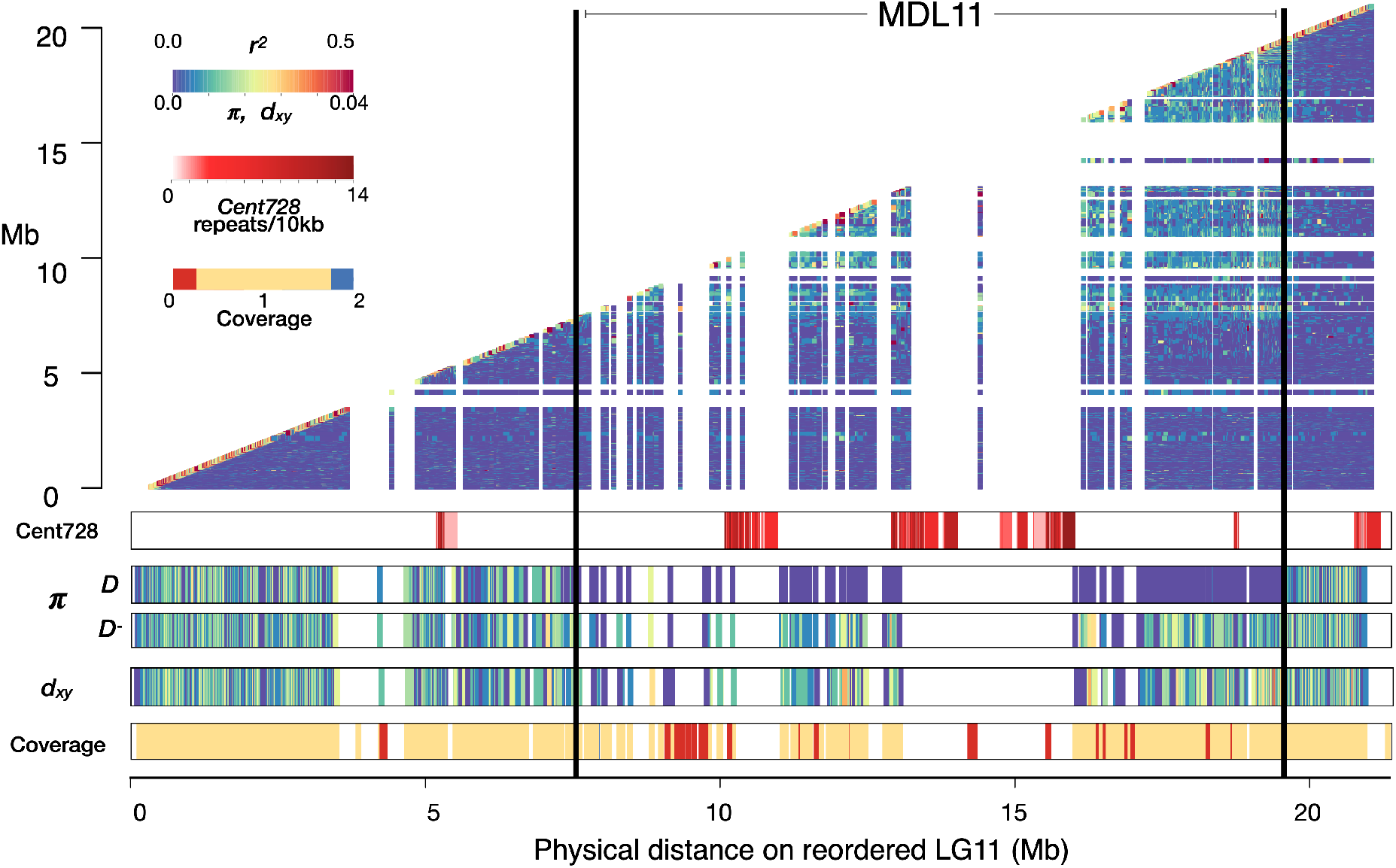
Elevated linkage disequilibrium (*r^2^*) and reduced diversity (π) define a distinct haplotype around expanded *Cent728* repeats associated with female meiotic drive in *Mimulus guttatus*. A heatmap of pairwise estimates of *r^2^*, plotted by megabases (Mb) on x- and y-axes, illustrates the region of suppressed recombination corresponding to Meiotic Drive Locus 11 (MDL11 in the Iron Mountain (IM) population of *M. guttatus* (N = 34 IM inbred lines). Lower panels (in order from top to bottom) show the chromosome-wide density of putatively centromeric Cent728 repeats, nucleotide diversity (π) per gene for lines carrying driving *D* (N = 14) and non-driving *D*^−^ haplotypes (N= 20), divergence (*d_x,y_*) per gene between D and *D*^−^ lines, and the ratio of coverage in *D*^−^ lines vs. *D* lines when both are aligned to the *D* reference genome (values near zero indicate likely deletion in *D*^−^ vs. *D*, whereas values near 2 indicate possible duplication).

As predicted by population genetic models (Fishman and Kelly 2015) and previously inferred from a handful of marker sequences (Fishman and Saunders 2008), the sweeping away of genetic variation demonstrates the rapid and recent spread of *D* to intermediate frequency despite substantial individual fitness costs. Across MDL11, *D* lines are essentially invariant whereas *D*^−^ lines are highly variable and both sets of lines exhibit high diversity in flanking regions (Table 1, Fig. 1, Fig. S1). To estimate the age of the recently swept *D* haplotype within the IM population, we counted single nucleotide variants (SNVs) in coding sequence across the region in 13 *D* lines (Supporting Information; Table S3). Over ~256 kb of unambiguously *D* coding sequence we identified 9 single nucleotide variants (SNVs) present in one or more lines. Using mutation rates = 0.2 - 1.5 × 10^−8^, following (Brandvain et al. 2014), this accumulation of variation corresponds to 200 - 1497 years (generations) since the sweep with simple population genetic equations (Thomson et al. 2000). Forward simulations with similar parameters find a mean time to common *D* ancestor of 999 years (Supporting Information; Fig. S2).

**Table 1.** Nucleotide diversity across LG11 in the IM *Mimulus guttatus* population.

| Lines<sup>a</sup> | Region<sup>b</sup> | Mean $\pi$ (SE)<sup>c</sup> | Mean $d_{x,y}$ (SE) | 95% CI<sup>d</sup> | N<sub>genes</sub><sup>e</sup> |
| --- | --- | --- | --- | --- | --- |
| *D* | MDL11 | 0.0002 (0.00009) | -- | (0.00007 - 0.0004) | 219 |
| *D*<sup>-</sup> | MDL11 | 0.0097 (0.0004) | -- | (0.0089 - 0.0104) | 219 |
| *D* | Flanking | 0.0096 (0.0002) | -- | (0.0092 - 0.0100) | 855 |
| *D*<sup>-</sup> | Flanking | 0.0100 (0.0002) | -- | (0.0096 - 0.0103) | 855 |
| *D* vs *D*<sup>-</sup> | MDL11 | -- | 0.0110 (0.0005) | (0.0102 - 0.0120) | 231 |
| *D* vs *D*<sup>-</sup> | Flanking | -- | 0.0098 (0.0002) | (0.0094 - 0.0101) | 867 |

<sup>a</sup>14 IM lines with *D* haplotype, 20 IM lines with *D*<sup>-</sup> haplotype
<sup>b</sup> MDL11 = region of LG11 spanning driving *D* haplotype; Flanking = LG11 outside of MDL11
<sup>c</sup>Nei's diversity per gene per site (Nei and Li 1979)
<sup>d</sup> Confidence intervals (CI) generated by resampling the mean without assuming normality (N = 1000)
<sup>e</sup>Number of genes without missing data

Given the distinctiveness of the *D* haplotype, it is worth considering whether it arose by local mutation, gene flow from another population, or introgression from another species. The *D* haplotype also occurs in at least one other intensively sampled population from the Oregon Cascades (Monnahan and Kelly 2017), suggesting that it may not have arisen by mutation within our focal population. However, both divergence estimates and coalescent models suggest that haplotype associated with drive is unusually extended and common, but not unusual in sequence or origin. Divergence (genic *d_x,y_*) between *D* and *D*^−^ lines is only marginally higher in the MDL11 region vs. flanking regions (0.011 vs. 0.0098; Table 1, Fig. 1). Further, while trans-specific introgression of other loci has been observed at Iron Mountain (Puzey et al. 2017), it is unlikely to be an initiator of drive in *M. guttatus.* In pairwise coalescent analyses with samples from outside the IM population, the *D* and *D*^−^ haplotypes exhibit similar inferred demographic histories, both inside and outside MDL11 (Fig. S3). Further, consistent with no elevation of *d_x,y_* across MDL11 (Table 1), there is no evidence of unusually deep coalescence between the sampled *D* and *D*^−^ haplotypes. Together, these results suggest that the driving *D* haplotype arose by structural and sequence mutation within the Northern clade of *M. guttatus* rather than from long-distance migration or interspecific introgression.

Given that the MDL11 region includes at least 387 protein-coding genes (Fig. 1, Table S4), it is possible that linked genes enhance female meiotic drive and/or contribute to the substantial costs of *D* homozygosity. Male meiotic drive factors, such as *Segregation Distorter* in fruit flies, are often associated with rearrangements that genetically link sperm-killing alleles with responder or enhancer genes (Larracuente and Presgraves 2012). Female meiotic drive, on the other hand, involves physical competition between structurally divergent chromosomes and thus does not require differences in gene sequence or expression. However, linked genic enhancers are predicted to accumulate whenever LD is high around any selfish element (Crow 1991). Furthermore, female meiotic drive by a neocentromeric driver in maize requires both a physical knob of heterochromatic satellite DNA and a cluster of kinesin genes, which are locked together within an inversion (Dawe et al. 2018). To assess the opportunity for collusion between driving *Cent728* arrays and linked genes, we surveyed MDL11 for genes with potential meiotic functions (Table S4). Candidates include the sole *M. guttatus* homologue of Nuclear Autoantigenic Sperm Protein (NASP)/Sim3, which was recently identified as the chaperone of plant centromeric histones (Le Goff et al. 2020). In addition, a > 800kb region (45 genes: Migut.K01214- Migut.K1259) exhibiting near-zero sequence coverage in all *D*^−^ lines Fig. 1, Table S4) contains a homologue of Arginine-Rich Cyclin RCY1, a component of the male-meiosis-essential Cyclin L/CDKG1 complex (Zheng et al. 2014). Thus, gene content differences between *D* and non-*D* haplotypes may also contribute to drive or its costs. However, because all diagnostic variants are equally associated with meiotic drive within the IM population and in hybrids, we cannot genetically uncouple these potential genic modifiers from the *Cent728* arrays. In the future, genetic editing of target sequences in *Mimulus* may make direct study of their drive-relevant functions possible.

Centromeric drive sets up a conflict between the driver and genes genome-wide, with components of the kinetochore machinery particularly likely evolutionary interactors. In *Mimulus guttatus*, the striking difference in the strength of drive between heterospecific and conspecific hybrids allows quantitative genetic investigation of this process over long time-scales, while costly drive polymorphism within IM can illuminate it from a population genetic perspective. Thus, we first ask whether unlinked suppressor loci contribute to the relative weakness of conspecific (*DD*^−^; 58:42) vs. heterospecific (*Dd;* 98:2) drive and then examine population genomic patterns of selection at a functional and positional candidate. These approaches are complementary: the quantitative genetic approach casts a broad net to assay accumulated differences between species but cannot distinguish driven co-evolution from other sources of epistasis in hybrids (Fishman and Sweigart 2018; Sweigart et al. 2019), while the population genomics is a single gene-scale snapshot of evolution in action.

Centromeric or genic divergence within MDL11 alone (i.e. *M. guttatus D*^−^ vs. *M. nasutus d* as competitors with *D*) could govern the strength of transmission ratio distortion in *DD*^−^ vs *Dd* heterozygotes. However, *M. nasutus* alleles at unlinked loci may be particularly permissive to drive in F_1_ hybrids and *M. nasutus*-background nearly isogenic lines (Fishman and Willis 2005). To evaluate these (non-exclusive) alternatives and map any unlinked modifier loci, we generated a three-parent interspecific F_2_ mapping population by crossing a *Dd* F_1_ female parent (SF *M. nasutus* x IM160 *M. guttatus*) to a *D-d* F_1_ male parent (SF x IM767), genotyped the F_2_ hybrids genome-wide using a reduced-representation sequencing method, and constructed a linkage map (Flagel et al. 2019). As expected, the *Dd* F_1_ female transmitted only *D* alleles to the next generation, and the F_2_ hybrids consisted entirely of *Dd* and *DD^−^* individuals (n = 88 and 96, respectively). We used the frequency of *D* in selfed-F_3_ progeny of F_2_ hybrids (n = 12-16 genotyped per family, total N > 2400) to calculate the strength of female meiotic drive (%D_fem_, assuming male to be Mendelian). Averaged across genetic backgrounds in F_2_ siblings, *Dd* drive remained dramatically stronger than *DD^−^* drive (mean %D_fem_ = 0.93 vs. 0.73; r^2^ = 0.26; n = 159). Thus, stronger drive against the *M. nasutus d* allele can primarily be ascribed to structural and/or genic divergence in the functionally centromeric MDL11 region. Because *M. nasutus* is a highly selfing species (Brandvain et al. 2014), centromeric drive and other forms of genetic conflict should have been relaxed since its split from *M. guttatus* (Burt and Trivers 1998). Thus, beyond the current dynamics of the *D* variant at MDL11, *M. nasutus* may have both generally “weak” centromeres and a cellular machinery that is particularly vulnerable to selfish elements.

Despite its predominant effect, genotype at MDL11 could not fully explain variation in the intensity of drive, suggesting that unlinked genetic modifiers also modulate drive in interspecific F_2_ hybrids. In our F_2_s, *Dd* drive (0.93) was reduced relative to the expectation from F_1_s and majority-*M. nasutus* isogenic lines (0.98) (Fishman and Willis 2005), whereas *DD*^−^ drive was substantially elevated relative to our expectation from previous crosses within IM (mean %*D*_fem_ = 0.73 vs. 0.58) (Fishman and Saunders 2008). A scan for quantitative trait loci (QTLs) affecting *D* transmission detected weak unlinked modifiers on LG9 and LG14 (LOD > 2.0; peak r^2^ = 0.09 for both; Fig. 2A). The large confidence intervals (20-50 cM) of these minor QTLs span hundreds of genes, but the LG14 modifier QTL is notably centered over one of the two genes encoding CenH3 in *M. guttatus* and relatives (CenH3A) (Finseth et al. 2015). Because CenH3 proteins are the leading functional candidates for suppression of centromeric drive (Henikoff et al. 2001), we further characterized *Dd* and *DD*^−^ drive in all four CenH3A genotypes found in our F_2_ hybrids (G_160_G_767_, NG_160_, NG_767_, and NN as determined by diagnostic marker alleles; N = 150). We see a strong primary effect of MDL11 genotype (F_1,3_ = 47.20, P < 0.00001) and (marginally) the expected elevation of *D* transmission in *M. nasutus* vs. *M. guttatus* CenH3A homozygotes across both MDL11 genotypic classes (Least Squares Means comparison: *P* = 0.059; Fig. 2B). In addition, CenH3A and MDL11 genotypes interacted non-additively (*F* = 3.91, *P* < 0.01), with *DD*^−^ drive becoming as strong as *Dd* drive exclusively in NG_767_ heterozygotes (Fig. 2B). Although the CenH3A allele from IM767 does not enhance conspecific drive when paired with a second *M. guttatus* allele, as in (Scoville et al. 2009), this allelic interaction intriguingly mirrors transgenic experiments transferring CenH3s among widely divergent plant species. In that work, *Arabidopsis* plants expressing homozygous maize CenH3 produce viable offspring when selfed, but engineered maize-*Arabidopsis* CenH3 heterozygotes exhibit zygotic mis-segregation, aneuploidy, and inviability (Maheshwari et al. 2015). This apparent underdominance implies uniquely negative interactions between distinct versions of CenH3 during cell division. Similarly, our results suggest that sensitivity of meiosis to mismatch between (some) heterospecific CenH3 alleles, on top of the posited role for mismatch between CenH3 and centromeric DNA (Henikoff et al. 2001), may unmask drive or cause reproductive breakdown in hybrids.

**Fig. 2.**
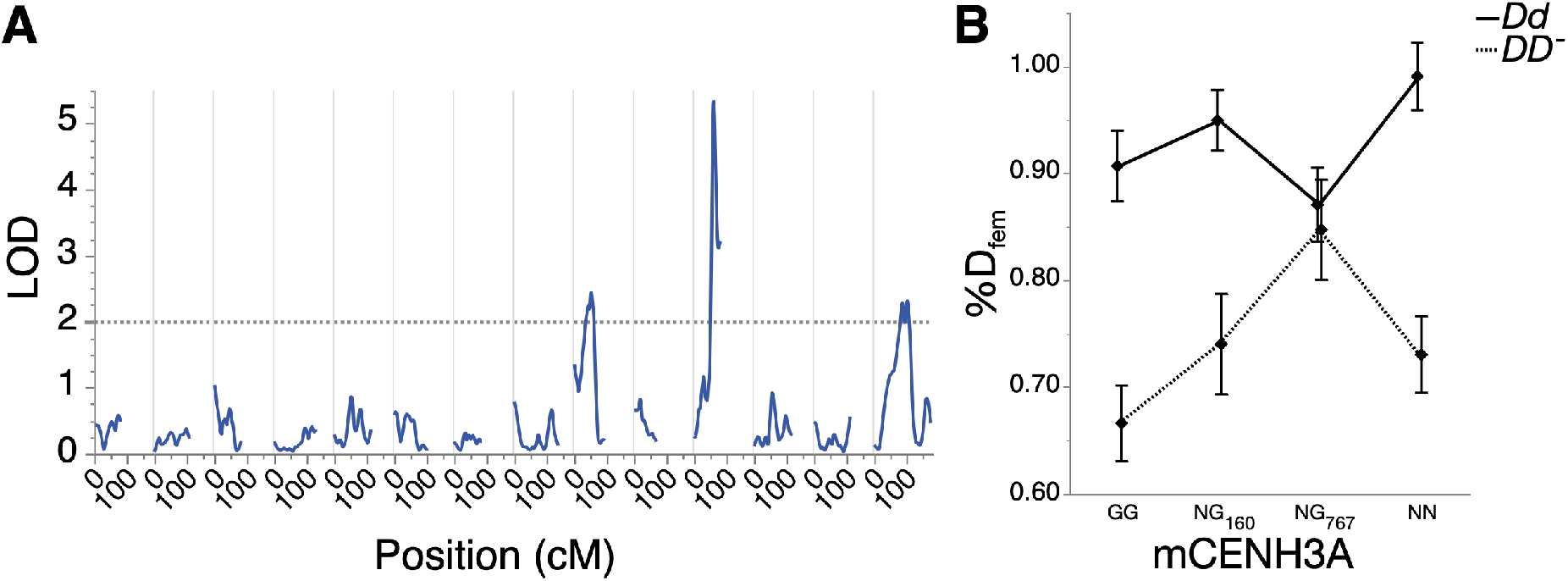
The relative strength of conspecific and heterospecific drive depends on the non-driving genotype at MDL11, as well as unlinked modifiers. **A.** A quantitative trait locus (QTL) scan of transmission ratio distortion in progeny of F_2_ hybrids reveals unlinked modifier QTLs on Linkage Groups/Chromosomes (LG) 9 and 14, in addition to the primary effect of MDL11 genotype. **B**. F_2_ genotype at CenH3A, which is centered under the LG14 modifier QTL, significantly influences *D* transmission in hybrids. Due to the three-parent crossing scheme (see Methods), there are only two F_2_ hybrid genotypes (*DD*^−^ and *Dd*) at MDL11, but four possible CenH3A genotypes: GG (IM160/IM767 *M. guttatus*), NG_160_ (heterozygote with *M. guttatus* allele from IM160 parent), NG_767_ (heterozygote with *M. guttatus* allele from IM160 parent), and NN (M. *nasutus*).

While quantitative genetic modification of drive by linked and unlinked genes in *M. nasutus* x M*. guttatus* hybrids likely reflects evolution in both species, the spread of *D* (with its costs) specifically predicts signatures of recent selection on interacting loci within the Iron Mountain *M. guttatus* population. We examined the two centromeric histones, as they are primary functional candidates for antagonistic co-evolution with costly *D* chromosomes and CenH3A is a candidate modifier in the mapping. Strikingly, an 8-gene region centered on CenH3A (Migut.N01552-Migut.N01559) exhibits a near-complete selective sweep at IM (Fig. 3, Fig. S4), whereas CenH3B shows no signatures of local selection (Finseth et al. 2015; Puzey et al. 2017). The CenH3A region is an outlier in within-population nucleotide diversity (mean π: 0.00232, *P* < 0.017) and has a significantly skewed site frequency spectrum (mean Tajima’s D: −0.838, P < 0.017, Fig. S4B), but exhibits typical inter-population diversity (*P* > 0.05 in all comparisons; Table S5). These signatures, along with elevated linkage disequilibrium (Fig. 3), indicate a recent local selective sweep rather than widespread purifying selection.

**Fig. 3.**
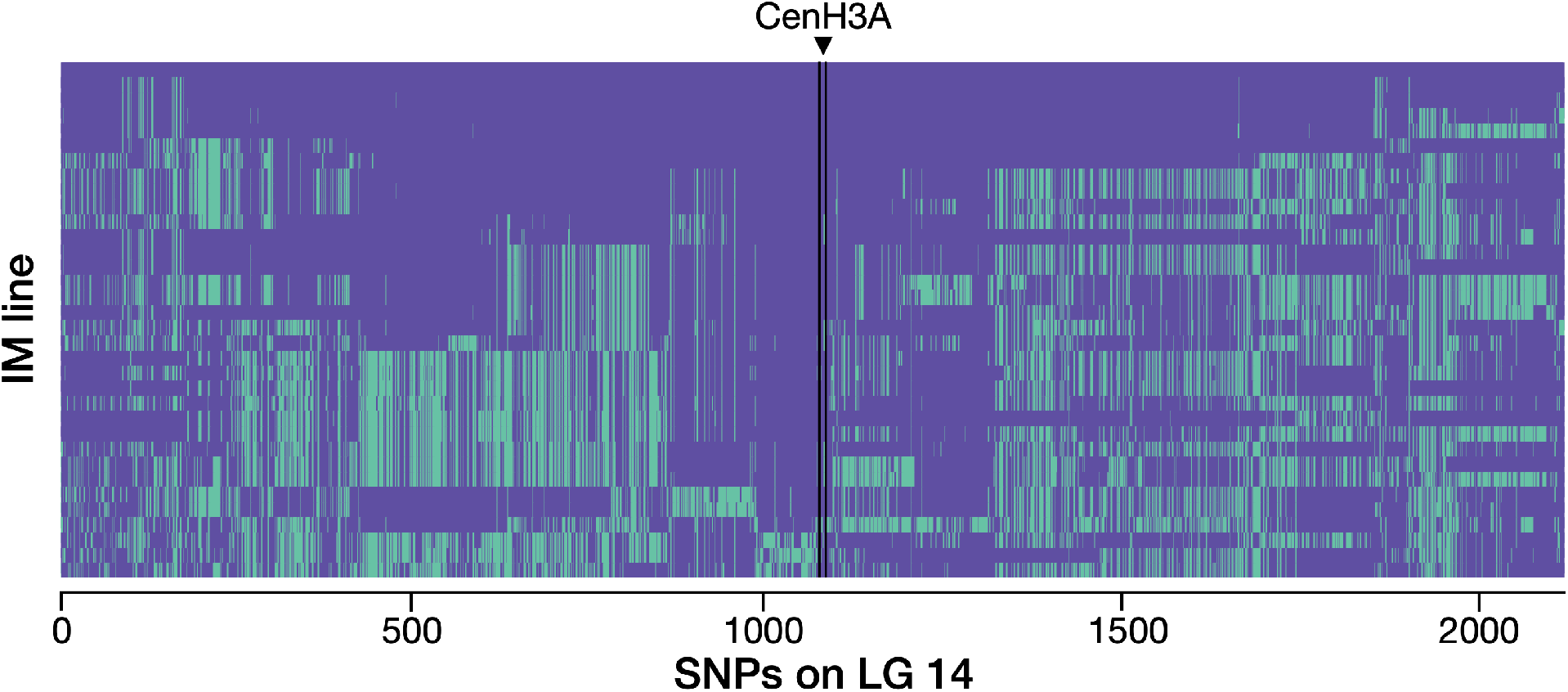
The genomic region around CenH3A exhibits a recent selective sweep in the Iron Mountain *Mimulus guttatus* population, consistent with evolution in response to the spread of costly *D* chromosomes. For each of 34 IM lines, single nucleotide polymorphisms (SNPs) across a 496 kilobase (kb) region flanking CenH3A are coded according to whether they match (purple) or differ (green) from IM1054, which bears one of the most common CenH3A-flanking haplotypes. The arrowhead and lines mark the location of CenH3A. The seven haplotypes were assigned manually and are detailed in Table S6; for visual resolution around CenH3A, the longest haplotype (> 620kb) was truncated.

To age the CenH3A selective sweep relative to that of *D*, we considered two scenarios. First, if the 23.9 kb low-diversity region immediately around the gene decayed (from single whole-chromosome haplotype) following the introduction of a novel mutation now near fixation, the selective sweep at IM occurred 627-4178 years ago, depending on the local recombination rate (Methods). However, strong haplotype structure extending across a substantially larger flanking region around CenH3A (Fig. 3) suggests that novel selection likely favored an existing variant found on multiple genetic backgrounds. Seven distinct long-range haplotypes of CenH3A were represented by two or more lines (Table S6), and the length of the two most common haplotypes (168.5 and 254.4 kilobases, respectively) supports a more recent response to novel selection (range 53 - 722 years ago; Methods). Of course, the history of selection on CenH3A may be far more complex than either of these scenarios. CenH3 sequences routinely exhibit the recurrent positive selection detected by measures of long-term molecular evolution (Finseth et al. 2015), and *D* may not be the only selfish centromere exerting selection in *M. guttatus* (or even at Iron Mountain). Nonetheless, the timescale of either scenario is consistent with the hypothesis that the recent spread of *D* to intermediate frequency (with attendant or subsequent fertility costs) sparked selection on CenH3A variation.

## DISCUSSION

Overall, our results demonstrate that genic factors can modify the strength of centromeric drive and that the recent spread of a selfish chromosome has plausibly driven local evolution of a key kinetochore protein in a wild plant. This supports models in which centromeres routinely drive through asymmetric female meiosis, with fitness consequences that select for compensation by other components of the segregation machinery. However, we note that all three steps of the centromere drive model occur simultaneously in our focal population of *Mimulus guttatus* only because *D* carries recessive costs that generate balancing selection (Fishman and Kelly 2015; Hall and Dawe 2018) and thus provide time for the rise and spread to fixation of even weak suppressor mutations (Crow 1991). This point is underlined by the finding that the CenH3A selective sweep at IM likely involved a standing variant found on multiple haplotypic backgrounds, and thus did not require that a modifier mutation fortuitously hit the small sequence target of core meiotic proteins. Although recessive costs of drive may be a direct consequence of meiotic conflict between driving centromeres, rearrangements that suppress recombination around them may slow the dynamics by creating opportunities for both costly hitchhikers and linked enhancers. Thus, centromeric drive and kinetochore protein coevolution, and their consequences for individuals, populations, and species, may be often be intertwined with the processes that shape the evolution of chromosome structure more broadly.

## METHODS

### Genome sequencing, alignment, read mapping, and data filtering protocols

Whole genome re-sequence data (fastqs, Illumina reads) were obtained from the Sequence Read Archive (SRA) for 34 Iron Mountain (IM) inbred lines and four lines (AHQT, DUN, LMC24, and MAR3) from distant populations (Flagel et al. 2014; Lee et al. 2016; Puzey et al. 2017). We generated new sequence data for two additional plants (one inbred line, one F_1_) derived from Iron Mountain. For the newly sequenced lines, DNA was extracted from fresh tissue using a modified CTAB-chloroform extraction protocol dx.doi.org/10.17504/protocols.io.bgv6jw9e. New genomic libraries were prepared following the Nextera tagmentation protocol and sequenced on the Illumina NextSeq platform (Ilumina NextSeq paired-end, 150 bp reads; Ilumina Inc., San Diego, USA), as described in (Case et al. 2016). All samples and their populations of origin, MDL11 haplotype call, and source are detailed in Table S2. Note that IM712 was only included in linkage disequilibrium (LD) and depth of coverage analyses.

All sequences were quality- and adapter-trimmed with Trimmomatic version 0.35 (Bolger et al. 2014) and aligned to the *M. guttatus* v2 reference genome (www.Phytozome.jgi.doe.gov) using bwa mem version 0.7.15 with default parameters (Li and Durbin 2009). Reads with mapping qualities less than 29 were filtered out with SAMtools v 1.3 (Li et al. 2009) and duplicate reads were removed (Picard tools v 1.119; http://broadinstitute.github.io/picard). We used the Genome Analysis Toolkit (GATK) to re-align around indels and call variant sites with the Unified Genotyper tool, following GATK’s Best Practice recommendations (McKenna et al. 2010; DePristo et al. 2011). Datasets were restricted to bi-allelic positions within genes using vcftools v0.1.12b (Danecek et al. 2011), indels were removed, and sites covered by less than three reads per line were converted to missing data. For the highly inbred IM lines (mean H_OBS_ per individual = 0.041, SD = 0.01), we removed sites with any heterozygous genotype calls. For population genomic analyses, sites with genotype calls from at least 10 individuals were retained and genes with fewer than 150 retained sites were removed. Comparisons between IM and lines from distant populations (AHQT, DUN, LMC24, and MAR3) were restricted to sites retained in the IM population.

### Characterization of the MDL11 region

#### Scaffold re-ordering

For analyses of sequence variation on LG11, we used a physical map based on the re-ordering of *M. guttatus* v1 scaffolds in a collinear (*D*^−^ × *D*^−^) IM767 x Point Reyes *M. guttatus* F_2_ mapping population (Holeski et al. 2014; Flagel et al. 2019). In this mapping, v1 scaffolds were re-positioned, split, and inverted to optimize the genetic map, while retaining sequence and gene-annotation information for each v1 segment from the v2 assembly. In addition, we included the large (> 3 Mb) gene-poor v1 scaffold_10 in the MDL11 region (Table S1), as it was placed there in v2 (and is clearly part of the *D* haplotype block in visual examination of Illumina-read alignments), but was lost from later genetic maps due to low genotyping quality in this repetitive region (Flagel et al. 2019). Mapped v1 scaffold sequences were extracted from the v2 reference genome and reordered into a new FASTA file based on their genetic coordinates. All gene sequences between contiguous genetically-mapped 100kb v1 segments were included in LD analyses (making them conservative; 1,188 genes), but divergent genes that were clearly not part of the MDL11 haplotype block (likely due to local mis-assembly) were excluded in remaining analyses unless specified (1,104 genes included; Table S4).

#### Localization of *Cent728* satellite repeats and analyses of gene content

We used the Basic Local Alignment Search Tool (BLAST) (Altschul et al. 1990) of the consensus nucleotide sequence of *Cent728* (Fishman and Saunders 2008) to localize copies of the putative centromeric repeat on the re-ordered LG11. To survey for gene content differences (copy number variation; CNV) between *D* and *D*^−^ individuals across LG11, we used deviations in read depth following (Nelson et al. 2018). We allowed sites to have missing data and relaxed the read coverage per line criteria for these analyses. We used vcftools v0.1.12b (Danecek et al. 2011) to obtain read depth for each exonic site (excluding indels, heterozygous sites, and sites with more than two alleles), standardized values by the individual’s chromosome-wide median for such sites, and calculated an average standardized read depth for each gene. Genes were excluded as likely misassembled or repetitive if *D* individuals had standardized coverage values < 0.5 or > 3, or if they were identified as chloroplast-nuclear transfers or nongenic mis-annotations in (Nelson et al. 2018). On LG11, 1,344 genes were retained. *D^−^:D* coverage ratios were used to categorize genes as likely deleted (0 – 0.25; red), duplicated (1.75 – 2.0; blue), or not likely duplicated or deleted (0.25 - 1.75; tan; Fig. 1).

#### Linkage disequilibrium, nucleotide diversity, and site frequency spectrum

To estimate linkage disequilibrium across LG11, we used vcftools version 1.12a (Danecek et al. 2011) to calculate the squared correlation coefficient between genotypes (r^2^) for SNP pairs (N = 49,595 genic SNPs at IM). Average r^2^ across all polymorphic sites was then calculated for each gene pair (N = 1,475 genes). Second, we explored haplotype structure by calculating the proportion of SNPs per gene on LG11 that matched the IM62 reference for each sequenced line. For the haplotype structure analyses, we coded genes with fewer than seven polymorphic sites genotyped as missing data (N =1,064 genes included). Average within-population nucleotide diversity (π) per gene, as well as *d_x,y_* between *D* and *D*^−^ lines, was calculated in R using PopGenome (Pfeifer et al. 2014), based on (Nei and Li 1979), and divided by the number of sites per gene. Calculations were performed separately for IM lines with *D* and *D*^−^ haplotypes, and values were averaged over MDL11 and flanking regions, respectively, in each. Genes inside MDL11 are listed in Table S4; all other genes were considered to be in flanking regions. Confidence intervals were generated in the Hmisc package of R, version 4.1-1, by performing 1000 bootstrap re-samplings of the means without replacement (Harrell 2018).

#### Origin and age

To infer the demographic history of the MDL11 region, we applied pairwise sequentially Markovian coalescent (PSMC) analyses as implemented by (Li and Durbin 2011). Following (Brandvain et al. 2014), we created pseudo-diploids by combining haploid genomes from two inbred lines for estimation of pairwise coalescence and effective population size through time. To place *D* in context, we used two non-IM *D*^−^ lines with distinct evolutionary affinities: a coastal perennial individual derived from the Southern *M. guttatus* clade (DUN) and an annual representing the Northern *M. guttatus* clade (MAR), to which IM also belongs (Brandvain et al. 2014), as well as *D* (IM62) and *D*^−^ (IM767) IM lines. For this analysis, bams were first made as described in (Case et al. 2016). Pseudo-diploids were then created by making fasta files using the consensus sequence of each bam and merging the two consensus sequences using the seqtk toolset (https://github.com/lh3/seqtk) with a quality threshold of 20. We performed 100 bootstrap replicates for each pairwise comparison. To perform the bootstraps, we applied the splitfa tool from the PSMC package to break the pseudo-diploids into non-overlapping chunks. The segmented genome then served as input for 100 separate PSMC analyses with the -b option. Coalescent analyses were performed separately for chromosomal locations within MDL11 and in flanking regions (Table S4).

Because *D* is non-recombining with alternative alleles, we used mutation alone (rather than haplotype structure or a mix) to age it. First, to estimate the time since most recent common ancestor (t) for the *D* haplotype, we counted the number of segregating sites in 13 IM lines (IM62, IM115, IM116, IM138, IM1145, IM239, IM502, IM657, IM664, IM742, IM909, IM922, IM1054, excluding IM549 due to low coverage; Table S3). We restricted this analysis to exonic sites where alignments are more reliable (Puzey et al. 2017). We excluded heterozygous sites and entire genes with >5 heterozygous exon sites, as these generally represent stacked copy number variants or other instances of incorrect alignment, which can also produce (apparently) homozygous SNPs.

To estimate the age of the swept *D* haplotype, we used both simple calculations (the Thomson estimator; Thomson et al. 2000)) and forward simulations using a range of mutation rates (0.2 × 10^−8^ - 1.5 × 10^−8^), following(Brandvain et al. 2014). The Thomson estimator tends to underestimate time to the most recent common ancestor, as it does not include the initial spread of the focal haplotype to high frequency (Thomson et al. 2000); however, this is not a major concern given the short time to equilibrium frequency expected for driving *D* (Fishman and Kelly 2015). We also simulated mutation accumulation on a nonrecombining chromosome using the simulation software SLiM 2.6 The D haplotype contains a total of 256,867 nucleotide positions for which we have high-confidence genotype calls. We therefore simulated a population of nonrecombining chromosomes of length 256,867 bp that begins as a small founder population (n = 20 chromosomes) and grows exponentially by 10% per generation to a stable size of either 50,000 chromosomes (89 generations of growth) or 500,000 chromosomes (113 generations). We sampled 13 chromosomes per generation and counted the number of observed segregating sites in the sample. Simulations ended and the generation number was recorded when 9 segregating sites were observed. We performed simulations over a range of per-base mutation rates to correspond to population-scaled mutation rates (4Nμ) of 0.001 (1 × 10^−8^ per generation for 50,000 chromosomes, 1×10^−9^ for 500,000 chromosomes), which is well below observed π at Iron Mountain, to 0.01, which is similar to π at IM (Puzey et al. 2017). Simulation results are plotted in Fig. S2.

### Genetic mapping of loci underlying interspecific differences in vulnerability to *D* drive

#### Crossing design

To test for unlinked modifiers of LG11 *D* drive, we inter-crossed heterospecific *Dd* (SF *M. nasutus* x IM160) and *D-d* (SF *M. nasutus* x IM767) F_1_ hybrids to form an F_2_ mapping population. Because the SF x IM160 F_1_ (*Dd*) was used as the female parent, we expected these F_2_s to all be *D*d or *DD*^−^ (no *dd*, due to near-complete drive in the female *Dd* parent). Thus, we can examine the strength of heterospecific (*Dd*) and conspecific (*DD*^−^) drive in a segregating F_2_ background and map any major loci that modulate their expression. F_2_ individuals were grown in a greenhouse at the University of Montana under standard long-day growth conditions for *M. guttatus*, and DNA was extracted from leaf tissue for genotyping using our standard 96-well CTAB-chloroform protocol (dx.doi.org/10.17504/protocols.io.bgv6jw9e). We then categorized individuals as *DD*^−^ (conspecific drive heterozygote) or *Dd* (heterospecific drive heterozygote) using the diagnostic marker Lb5a (Fishman and Saunders 2008).

#### Phenotyping

To characterize the strength of drive (the phenotype) in F_2_s, we hand self-pollinated 1-5 flowers per individual and collected the resultant selfed seeds. Some F_2_ hybrids set no seed, in part due to the segregation of known hybrid sterility factors (Sweigart et al. 2006) in this cross. For each selfed F_3_ seed family, we then planted 16 cells of a 96 well flat with 2 seeds each (or fewer if we did not have 32 viable seeds), and then thinned (and/or transplanted) to 16 per family. F_3_ plants were harvested as rosettes for DNA extraction and genotyping at diagnostic markers. Overall, we planted 250 progenies, and obtained 221 families (*Dd*; n = 101, *DD^−^;* n = 120) with at least 8 progeny successfully genotyped. For each progeny set, we estimated the strength of female meiotic drive (%D_fem_), assuming no distortion through male function (*Dd* expected > 0.98, *DD*^−^ expected = 0.58). This approach is not as precise as isolating female meiotic drive by hand-backcrossing F_2_s as dams (with prior emasculation in the bud) (Fishman and Willis 2005; Fishman and Saunders 2008), but selfing was more tractable for the large number of small-flowered F_2_s involved.

#### Linkage and quantitative trait locus (QTL) mapping

We constructed a linkage map of the F_2_ population (n = 184 total genotyped; 91 included in linkage mapping set) using multiplex shotgun genotyping (MSG) to generate low-coverage genome sequence (Andolfatto et al. 2011). The GOOGA pipeline (Flagel et al. 2019) was used to assign genotype probabilities to 100kb windows of the *M. guttatus* reference genome (v1 scaffolds; www.Phytozome.jgi.doe.gov) and order them into linkage groups corresponding to the 14 chromosomes of the *M. guttatus* and *M nasutus* genomes, as well as previous linkage maps of this interspecific cross (Fishman et al. 2001; 2014). As previously described (Flagel et al. 2019), this approach corrects numerous ordering errors in the v2 chromosome-scale assembly of *M. guttatus,* while also allowing use of the v2 annotation through assignment of each 100kb v1 segment to its corresponding v2 segment. This process resulted in 1,836 physically and genetically mapped window-based markers.

For QTL mapping, we used the posterior probabilities generated by GOOGA (Flagel et al. 2019) to make hard genotype calls for each 100kb genome window. Windows were assigned to one of the three fully informative genotypes (*M. guttatus* homozygote, *M. nasutus* homozygote, or heterozygote) if that genotype had a probability > 0.8. Windows that did not meet this criterion were called as missing. To verify that our genome-wide genotyping approach was effective, we tested for concordance between MDL11 windows and our D-diagnostic marker, excluding several individuals (likely contaminated during the MSG protocol and/or low coverage) where genotypes did not match. For QTL mapping of potential modifier loci, we restricted analyses to F_2_ individuals whose value of %D_fem_ was based on 12 or more F_3_ progeny, and who had <50% missing data (most much lower; N = 130). We scanned for QTLs underlying %D_fem_ using the interval mapping function in WinQTLCart (Wang et al. 2005), with marker-based D genotype as a binary co-factor. We used a generous significance threshold of LOD = 2.5 (p < 0.05) for the initial scan.

To characterize the LG14 modifier QTL, we made an exon-primed marker (mCenH3A; Table S7) that identified all three parental alleles of CenH3A – N from SF M. *nasutus*, G_160_ from IM160 and G_767_ from IM767 – based on length polymorphisms generated by intronic insertions and deletions. The two IM *M. guttatus* alleles were distinguished by a 1 basepair indel in the second intron. Because the crossing work pre-dated the sequencing of many inbred IM lines and the IM160 line was later lost, only an IM160 x IM767 F_1_ individual was available to sequence (Table S2). However, it is apparent that the IM160 allele of CenH3A happened to be unusually divergent, with >22 Single Nucleotide Polymorphisms (SNPs) and/or indels in introns and UTRs, one synonymous SNP in Exon 1, and one nonsynonymous SNP in Exon 4 (part of the rapidly evolving N-terminal tail) relative to both the reference and IM767. We genotyped mCenH3A in 150 F_2_s with >12 progeny contributing to their %D_fem_ phenotype, and tested for effects of the four possible genotypes (NN, NG_160_, NG_767_, and G_160_ G_767_) using a two-way analysis of variance with mCENH3A genotype, MDL11 genotype, and their interaction as factors (SAS Institute 2018).

### Population genomics of CenH3A

Average pairwise nucleotide diversity (π) per site per gene and Tajima’s D per gene were calculated for genes on LG14 (N = 2703) in R using PopGenome (Pfeifer et al. 2014), with the same parameters as for the analyses of LG11. CENH3A (Migut.N01557) resides in an 8-gene region of low diversity (Migut.N01552 – N01559; π < 0.005 in IM), which was also one of only 41 windows containing monophyletic-within-IM outliers in a previous study of selective sweeps at IM (Puzey et al. 2017). To further test whether such an extensive block of diversity reduction was extreme, we conducted permutations (N = 500) by calculating mean π for randomly chosen contiguous blocks of 8 genes along LG14. Confidence intervals were generated in the Hmisc package of R, version 4.0-2, by performing 1000 bootstrap re-samplings of the means without replacement (Harrell 2018).

To test whether diversity reduction around CenH3A at IM reflected low overall diversity, we also computed nucleotide diversity between samples from IM and distant populations, using the same approach as above. Calculations were performed sequentially between all IM lines and one other line (AQHT, DUN, MAR3, and LMC24), and confidence intervals generated as described above.

We visualized haplotype structure surrounding CenH3A using R version 3.5.0. Exonic SNPs on LG14 were phased using Beagle 4 (Browning and Browning 2007) and the haplotypes surrounding CenH3A (scaffold positions 13,500,000-14,000,000) were converted to a matrix using a custom Python script (vcf2selscan.py). We included one haplotype per inbred line and plotted allelic states at each SNP relative to the sIM1054 haplotype in R. Haplotypes were identified manually and their lengths are detailed in Table S6.

To estimate the age of the CenH3A sweep from the length of surrounding haplotypes, we followed the approach of (Sweigart and Flagel 2015), using a range of local recombination rates (150kb-1000kb/cM based on LG14 genetic maps). Because we have a broad distribution of haplotype lengths, we calculated ages using the shortest shared core segment (24 kb), as well as the longest, shortest, and mean haplotype lengths (Table S6). The latter bookend the age of the shift in selection from 24 years (longest, least recombination) to 722 years (shortest, most recombination). Because we do not currently have resolution to more finely estimate intra-population recombination, the key variable, we did not forward simulate this apparent sweep from standing variation.

### Confirmation of *D* vs. *D*^−^ gene content differences

Coverage differences between *D* and *D*^−^ lines at IM indicate that a 45-gene region is a) deleted in *D*^−^ relative to the (ancestral) *D* reference, b) inserted in *D* relative to ancestral *D*^−^ or c) so divergent that few or no reads from the *D*^−^ haplotype map to the *D* reference. The third alternative is unlikely, as exonic reads from across the species complex and beyond map well to exonic sequences in the IM62 reference (Brandvain et al. 2014; Garner et al. 2016). To further rule out this possibility, we designed an exon-primed, intron-spanning, length polymorphic PCR marker in the RCY1 homolog Migut.K01228/ Migut.K01229 (mK1229; Table S7). This marker also amplifies a fragment from a second RCY1 gene on LG10 (Migut.J00575), which acts as a positive control for amplification of the sample. We genotyped 120 wild-derived greenhouse-grown IM outbred plants using touchdown PCR amplification of fluorescently-tagged fragments sized with capillary electrophoresis on an ABI 3130 Genetic Analyzer (Fishman and Willis 2005). A 173bp fragment from Migut.K01229 segregated as a presence/absence polymorphism in perfect association with our standard MDL11 diagnostic marker for the IM population (Lb5a), while the 180bp band from Migut.J00575 was present in all individuals. This pattern (along with the low coverage shown in Fig. 1) suggests that the *D*^−^ plants do indeed lack sequence in this region.

## ACKNOWLEDGMENTS

We thank A. L. Sweigart, J. K. Kelly, A. Kern, and P. Ralph for helpful discussions, A.L. Sweigart, D. Crowser, A. Stathos, and K. Anderson for assistance with data collection, and L. Flagel, J.K. Kelly, J. H. Willis and J. H. Puzey for sharing sequence datasets. Funding: Supported by NSF grants DEB-0846089, DEB-1457763, and OIA-1736249 to L.F. and by Keck Science Department start-up funds to F.R.F.

## AUTHOR CONTRIBUTIONS

F.R.F., T.C.N. and L. F. collected the data, conducted the analyses, and wrote the paper.

## COMPETING INTERESTS

Authors declare no competing interests.

## DATA AND MATERIALS AVAILABILITY

The sequence data for all lines analyzed are available at the NCBI Sequence Read Archive as accessions listed in Table S2. Genotype-phenotype data will be archived on Dryad.

## Notes

### Competing Interest Statement

The authors have declared no competing interest.

