## Supplemental Figures and Tables for "Selfish chromosomal drive shapes recent centromeric histone evolution in monkeyflowers"

**This file includes:**

Figures S1 to S4
Tables S1 - S7

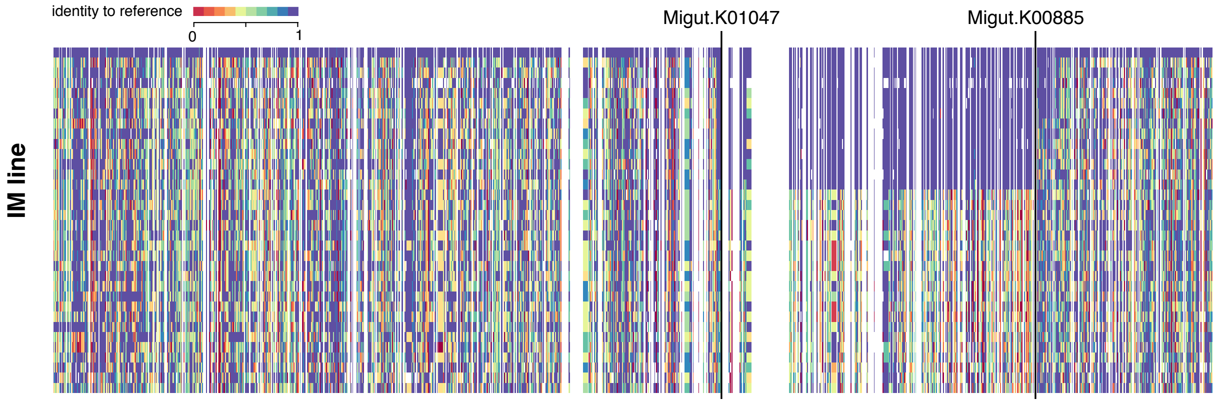


**Fig. S1.** The driving *D* haplotype of MDL11 is an extended region of sequence identity to the reference (IM62 line) *M. guttatus* genome, exhibited by ~1/3 of Iron Mountain *M. guttatus*. Thin black vertical lines bound the first and last gene of the MDL11 region (Table S4). Vertical bars are colored to represent proportion of SNPs in a gene that match IM62 (N = 1,064; genes with insufficient data are coded in white). Horizontal axis represents 34 individual inbred lines isolated from the IM population, from top to bottom: IM62, IM115, IM239, IM549, IM657, IM742, IM502, IM138, IM1054, IM922, IM909, IM664, IM116, IM1145, IM835, IM767, IM693, IM624, IM479, IM109, IM785, IM777, IM709, IM667, IM275, IM266, IM238, IM179, IM170, IM1192, IM1152, IM359, IM106, IM412.

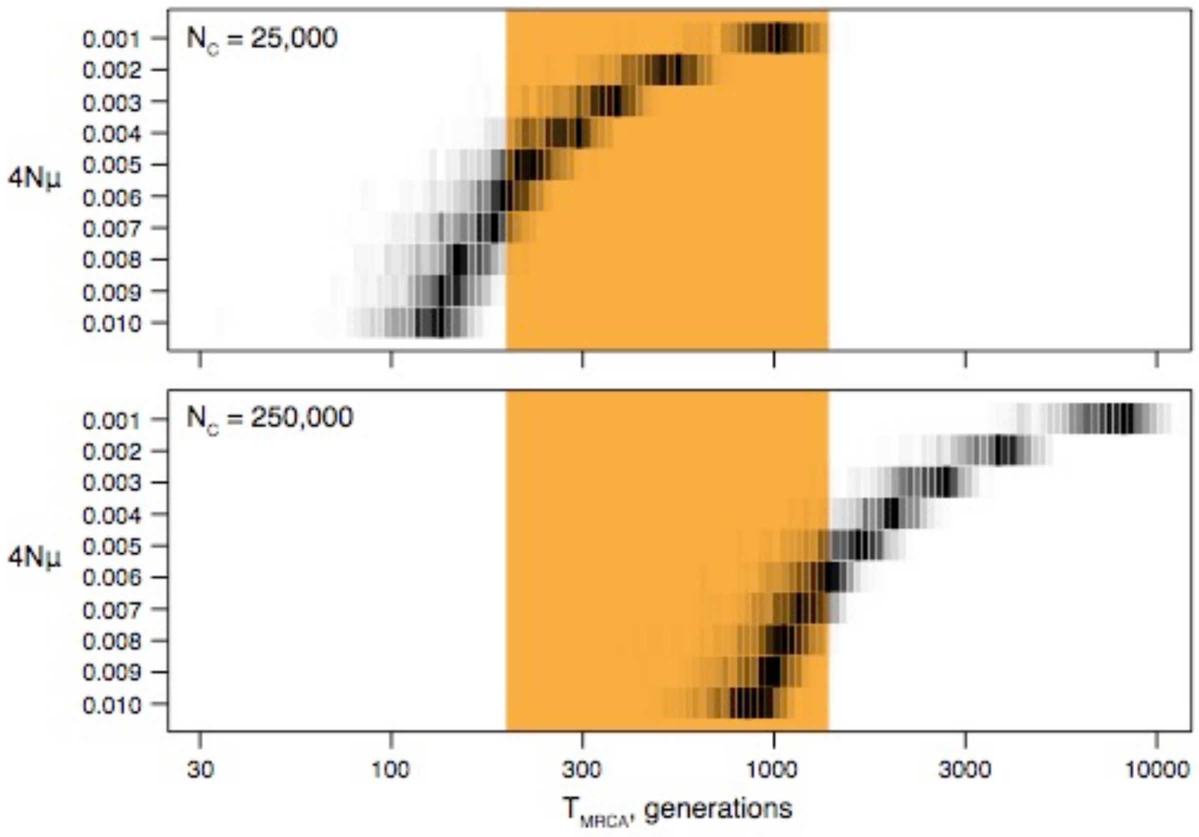


**Fig. S2.** Simulated and analytical results point to a recent origin of the *D* haplotype. Forward simulations were performed using SLiM 2 (described in Materials and Methods) over a range of mutation rates and with equilibrium census population sizes ($N_C$) of 25,000 (50,000 *D* chromosomes, top panel) and 250,000 diploids (bottom panel). Mutation rates are scaled by $N_C$ in the figure and correspond to ranges of $1x10^{-8}$-$1x10^{-7}$ (top) and $1x10^{-9}$-$1x10^{-8}$ (bottom). Grayscale density reflects the proportion of simulations yielding a $T_{MRCA}$ of the *D* haplotype within each bin. Gold shading represents the range of *D* haplotype ages calculated using the Thomson estimator (Thomson et al. 2000).

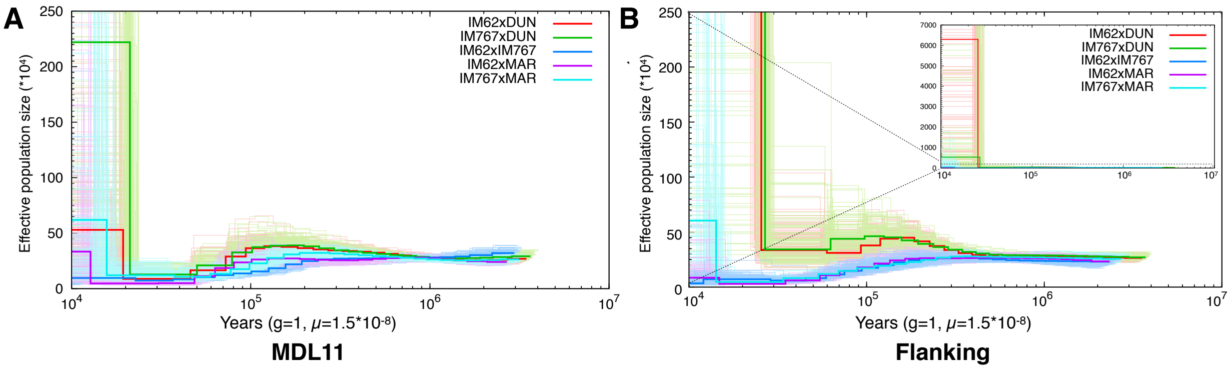


**Fig. S3.** The pairwise sequentially Markovian coalescent (PSMC) method suggests that *D* and $D^-$ alleles share a similar demographic history across LG11. PSMC inference of population size through time for pairwise haploid genome comparisons in the A) MDL11 region and B) the flanking regions of LG11. In B), the inset is a zoomed-out view of PSMC simulations. Haploid genomes of two inbred lines were used to create pseudo-diploids to use for estimating coalescence. Color codes are as follows: red = *D* line (IM62) x Southern-clade *M. guttatus* line (DUN); green = $D^-$ (IM767) line x southern *M. guttatus* (DUN); blue = *D* (IM62) line x $D^-$ (IM767) line; purple = *D* (IM62) line x northern *M. guttatus* (MAR); teal = $D^-$ (IM767) line x northern *M. guttatus* (MAR). Thick lines represent the point inference and thin lines represent bootstrap replicates (N = 100).

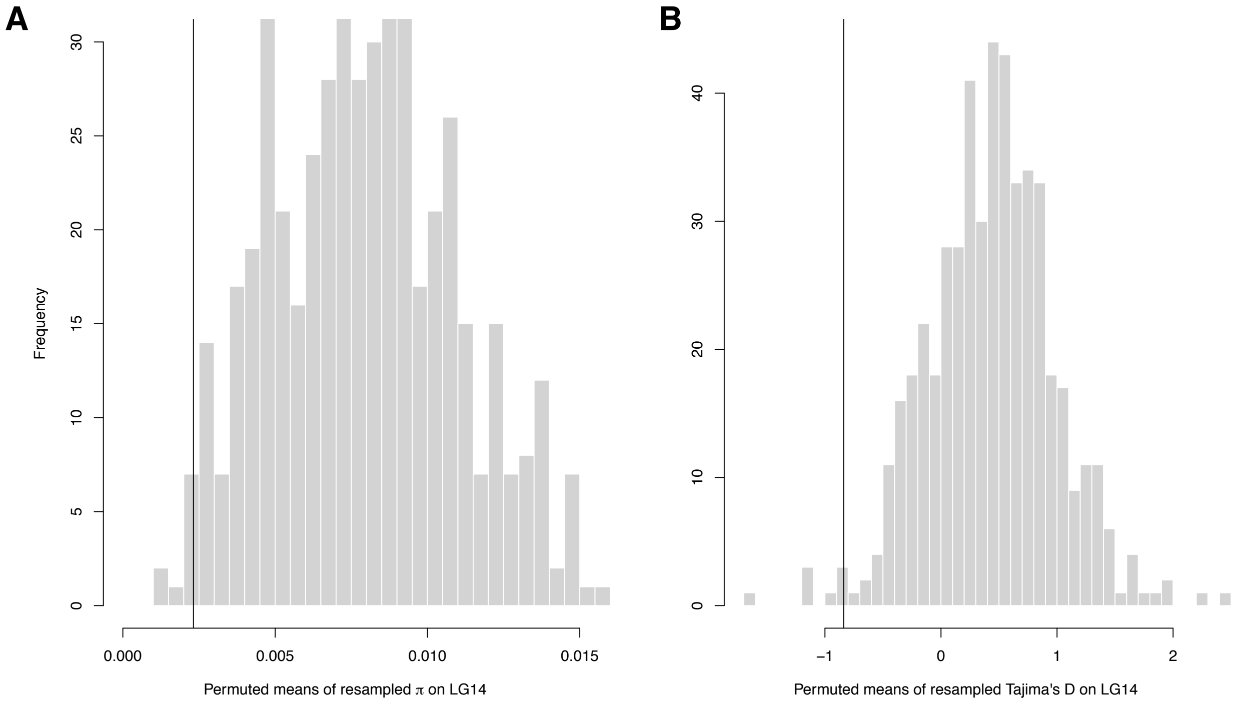


**Fig. S4.** Low nucleotide diversity (A) and a skewed site frequency spectrum (B) suggest that the 8-gene region around CenH3A (estimates indicated by vertical lines) experienced a selective sweep at IM (N = 34 lines). A) Histogram of permuted means calculated by averaging π per site per gene from blocks of 8 consecutive genes along LG14. Permutations were performed 500 times. B) Histogram of permuted means calculated by averaging Tajima's D per gene from blocks of 8 consecutive genes along LG14, which contains CenH3A. Permutations were performed 500 times.

Table S1: Re-ordered LG11 map. The order of LG11 based on a collinear $D^-$ x $D^-$ map (Flagel et al. 2019). For each *M. guttatus* v1 scaffold on LG11, the table shows assignment of its genes to *D* (1), $D^-$ (0) or both (REC), its v2 assembly position and genes, its orientation in the new map order, whether or not the v1 scaffold needed to be split, and its length.

| v1 scaffold | MDL 11? | v2 start | v2 end | Gene start | Gene end | v2 orient | v1 orient | v1 split | v1 length |
|---|---|---|---|---|---|---|---|---|---|
| 75 | 0 | 0 | 1159689 | K00001 | K00254 | F | F | N | 1159689 |
| 228 | 0 | 1169690 | 1556838 | K00255 | K00334 | F | F | N | 387148 |
| 273 | 0 | 1566839 | 1750548 | K00335 | K00367 | F | F | Y | 183709 |
| 213 | 0 | 1760548 | 2177483 | K00368 | K00445 | F | F | N | 416935 |
| 63 | 0 | 2430218 | 2187484 | K00485 | K00446 | R | R | Y | 242734 |
| 257 | 0 | 2605914 | 2440218 | K00513 | K00486 | R | F | Y | 165696 |
| 30 | 0 | 2615914 | 4575289 | K00514 | K00667 | F | R | N | 1959375 |
| 48 | 0 | 5742497 | 4831822 | K00736 | K00668 | R | F | Y | 910675 |
| 243 | 0 | 13523272 | 13853907 | K00948 | K00969 | F | F | N | 330635 |
| 415 | 0 | 14223853 | 14300664 | K00984 | K00987 | F | R | N | 76811 |
| 182 | 0 | 14427980 | 14963817 | K00994 | K01026 | F | F | N | 535837 |
| 185 | 0 | 15549682 | 15047665 | K01042 | K01032 | R | F | N | 502017 |
| 39 | REC | 16402937 | 15559683 | K01078 | K01043 | R | F | Y | 843254 |
| 47 | 1 | 18948653 | 17891937 | K01153 | K01123 | R | R | Y | 1056716 |
| 6 | 1 | 22410203 | 21063652 | K01260 | K01208 | R | F | Y | 1346551 |
| 145 | 1 | 11601986 | 12393390 | K00944 | K00944 | F | F | N | 791404 |
| 49 | 1 | 19506753 | 21053901 | K01154 | K01207 | F | R | N | 1547148 |
| 167 | 1 | 6237216 | 5673894 | K00751 | K00737 | R | R | N | 563322 |
| 10 | 1 | 6247217 | 9226930 | K00752 | K00759 | F | R | N | 2979713 |
| 239 | 1 | 13863908 | 14213852 | K00970 | K00983 | F | F | N | 349944 |
| 131 | 1 | 16412937 | 17176752 | K01079 | K01105 | F | R | N | 763815 |
| 221 | 1 | 17397001 | 17186753 | K01115 | K01106 | R | R | Y | 210248 |
| 100 | 1 | 11591985 | 10601358 | K00943 | K00886 | R | F | N | 990627 |
| 162 | 1 | 9900780 | 9236931 | K00809 | K00760 | R | F | N | 663849 |
| 161 | 1 | 10006245 | 10591357 | K00816 | K00885 | F | R | N | 585112 |
| 22 | REC | 22846476 | 24977148 | K01261 | K01499 | F | F | Y | 2130672 |

Table S2: Lines (inbred except for IM160xIM767 $F_1$) used in this study, with their population of origin, MDL11 haplotype, source and Sequence Read Archive accession number.

| Line | Location | MDL11 | Source | Accession # |
|---|---|---|---|---|
| DUN | Florence, OR | $D^-$ | Brandvain et al. 2014 | SRX030973 |
| IM1054 | Iron Mountain, OR | *D* | Puzey et al. 2017 | SAMN05852518 |
| IM106 | Iron Mountain, OR | $D^-$ | Puzey et al. 2017 | SAMN05852485 |
| IM109 | Iron Mountain, OR | $D^-$ | Flagel et al. 2014 | SRX021073 |
| IM1145 | Iron Mountain, OR | *D* | Flagel et al. 2014 | SRX021074 |
| IM115 | Iron Mountain, OR | *D* | Puzey et al. 2017 | SAMN05852486 |
| IM1152 | Iron Mountain, OR | $D^-$ | Puzey et al. 2017 | SAMN05852520 |
| IM116 | Iron Mountain, OR | *D* | Puzey et al. 2017 | SAMN05852487 |
| IM1192 | Iron Mountain, OR | $D^-$ | Puzey et al. 2017 | SAMN05852522 |
| IM138 | Iron Mountain, OR | *D* | Puzey et al. 2017 | SAMN05852490 |
| IM160 x IM767 | Iron Mountain, OR | $D \times D^- F_1$ | this study | PRJNA576504 |
| IM170 | Iron Mountain, OR | $D^-$ | Puzey et al. 2017 | SAMN05852491 |
| IM179 | Iron Mountain, OR | $D^-$ | Puzey et al. 2017 | SAMN05852492 |
| IM238 | Iron Mountain, OR | $D^-$ | Puzey et al. 2017 | SAMN05852495 |
| IM239 | Iron Mountain, OR | *D* | Puzey et al. 2017 | SAMN05852496 |
| IM266 | Iron Mountain, OR | $D^-$ | Puzey et al. 2017 | SAMN05852497 |
| IM275 | Iron Mountain, OR | $D^-$ | Puzey et al. 2017 | SAMN05852499 |
| IM359 | Iron Mountain, OR | $D^-$ | Puzey et al. 2017 | SAMN05852502 |
| IM412 | Iron Mountain, OR | $D^-$ | Puzey et al. 2017 | SAMN05852503 |
| IM479 | Iron Mountain, OR | $D^-$ | Flagel et al. 2014 | SRX021077 |
| IM502 | Iron Mountain, OR | *D* | Puzey et al. 2017 | SAMN05852504 |
| IM549 | Iron Mountain, OR | *D* | Puzey et al. 2017 | SAMN05852505 |
| IM62 | Iron Mountain, OR | *D* | Flagel et al. 2014 | SRX021072 |
| IM624 | Iron Mountain, OR | $D^-$ | Flagel et al. 2014 | SRX021075 |
| IM657 | Iron Mountain, OR | *D* | Puzey et al. 2017 | SAMN05852507 |
| IM664 | Iron Mountain, OR | *D* | Lee, Fishman et al. 2016 | SAMN04517335 |
| IM667 | Iron Mountain, OR | $D^-$ | Puzey et al. 2017 | SAMN05852509 |
| IM693 | Iron Mountain, OR | $D^-$ | Flagel et al. 2014 | SRX021078 |
| IM709 | Iron Mountain, OR | $D^-$ | Puzey et al. 2017 | SAMN05852510 |
| IM712 | Iron Mountain, OR | *D* | this study | PRJNA576504 |
| IM742 | Iron Mountain, OR | *D* | Puzey et al. 2017 | SAMN05852511 |
| IM767 | Iron Mountain, OR | $D^-$ | Flagel et al. 2014 | SRX021079 |
| IM777 | Iron Mountain, OR | $D^-$ | Puzey et al. 2017 | SAMN05852512 |
| IM785 | Iron Mountain, OR | $D^-$ | Puzey et al. 2017 | SAMN05852513 |
| IM835 | Iron Mountain, OR | $D^-$ | Flagel et al. 2014 | SRX021076 |
| IM909 | Iron Mountain, OR | *D* | Puzey et al. 2017 | SAMN05852516 |

| | | | | |
|---|---|---|---|---|
| IM922 | Iron Mountain, OR | *D* | Puzey et al. 2017 | SAMN05852517 |
| LMC24 | California | $D^-$ | Brandvain et al. 2014 | SRX030680 |
| MAR3 | Oregon | $D^-$ | Brandvain et al. 2014 | SRX030542 |
| AHQT | Wyoming | $D^-$ | Brandvain et al. 2014 | SRX142379 |

Table S3: Nine exonic mutations (by v2 Mb position) scored in 13 IM *D* lines, used for estimating the time since the *D* selective sweep. The ANC column contains the inferred sequence of the common ancestor of all sampled *D* haplotypes (found in six lines).

| Position (Mb) | ANC | 1054 | 115 | 116 | 138 | 664 | 239 | 502 | 657 | 742 | 909 | 922 | 1145 | 62 |
|---|---|---|---|---|---|---|---|---|---|---|---|---|---|---|
| 9413102 | G | G | G | G | G | G | G | G | G | G | G | G | **T** | G |
| 9420494 | T | T | T | T | T | T | T | T | T | T | T | T | **C** | T |
| 9664721 | G | G | G | G | G | G | G | G | G | G | G | **A** | G | G |
| 10228601 | A | A | A | A | **T** | A | A | A | A | A | A | A | A | A |
| 10739451 | A | A | A | A | A | A | **C** |  | A | A | A | A | A | A |
| 16858792 | T | T | T | T | T | T | T | T | T | T | T | **A** | T | T |
| 19962104 | G | G | G | G | G | G | G | G | G | G | G | G | **A** | G |
| 20024342 | C | C | C | C | C | C | C | C | C | **T** | C | C | C | C |
| 21424794 | A | **G** |  | A | A | A | A | A | A | A | A | A | A | **G** |

Table S4. Genes in the MDL11 region. Gene name, gene order in the reordered LG11 map, plus *Arabidopsis thaliana* best-hit homologue and gene descriptions from Phytozome 12.1.

| Migut # | Order | *Arabidopsis* homologue | Description |
|---|---|---|---|
| K01047 | 838 | AT1G11680 | CYTOCHROME P450 51G1 |
| K01046 | 839 | AT5G13400 | Major facilitator superfamily protein |
| K01045 | 840 | AT4G14100 | transferases, transferring glycosyl groups |
| K01044 | 841 | AT5G13390 | no exine formation 1 |
| K01043 | 842 | AT4G12420 | Cupredoxin superfamily protein |
| K01153 | 843 | | |
| K01152 | 844 | AT3G57680 | Peptidase S41 family protein |
| K01151 | 845 | | |
| K01150 | 846 | AT5G47900 | Protein of unknown function (DUF1624) |
| K01149 | 847 | AT3G51980 | ARM repeat superfamily protein |
| K01148 | 848 | AT5G47860 | Protein of unknown function (DUF1350) |
| K01147 | 849 | AT5G57330 | Galactose mutarotase-like superfamily protein |
| K01146 | 850 | | |
| K01145 | 851 | AT1G31260 | zinc transporter 10 precursor |
| K01144 | 852 | AT1G31260 | zinc transporter 10 precursor |
| K01143 | 853 | AT3G06240 | F-box family protein |
| K01142 | 854 | | |
| K01141 | 855 | AT1G31260 | zinc transporter 10 precursor |
| K01140 | 856 | | |
| K01139 | 857 | | |
| K01138 | 858 | AT1G67720 | Leucine-rich repeat protein kinase family protein |
| K01137 | 859 | AT4G23730 | Galactose mutarotase-like superfamily protein |
| K01136 | 860 | AT1G44960 | SNARE associated Golgi protein family |
| K01135 | 861 | AT1G28190 | |
| K01134 | 862 | AT1G44900 | minichromosome maintenance (MCM2/3/5) family protein |
| K01133 | 863 | AT1G21280 | |
| K01132 | 864 | AT5G47920 | |
| K01259 | 875 | AT4G29900 | autoinhibited Ca(2+)-ATPase 10 |
| K01258 | 876 | AT3G46920 | Protein kinase superfamily protein with octicosapeptide/Phox/Bem1p domain |
| K01257 | 877 | AT1G34070 | |
| K01256 | 878 | AT5G58200 | Calcineurin-like metallo-phosphoesterase superfamily protein |
| K01255 | 879 | AT3G21180 | autoinhibited Ca(2+)-ATPase 9 |
| K01254 | 880 | AT4G38950 | ATP binding microtubule motor family protein |
| K01253 | 881 | AT1G13750 | Purple acid phosphatases superfamily protein |

| | | | |
|---|---|---|---|
| K01252 | 882 | AT5G47770 | farnesyl diphosphate synthase 1 |
| K01251 | 883 | AT3G27110 | Peptidase family M48 family protein |
| K01250 | 884 | | |
| K01249 | 885 | AT3G02870 | Inositol monophosphatase family protein |
| K01248 | 886 | | |
| K01247 | 887 | | |
| K01246 | 888 | AT5G47770 | farnesyl diphosphate synthase 1 |
| K01245 | 889 | AT3G03730 | F-box family protein with a domain of unknown function (DUF295) |
| K01244 | 890 | | |
| K01243 | 891 | | |
| K01242 | 892 | | |
| K01241 | 893 | AT2G30360 | SOS3-interacting protein 4 |
| K01240 | 894 | | |
| K01239 | 895 | | |
| K01238 | 896 | AT3G24100 | Uncharacterised protein family SERF |
| K01237 | 897 | AT3G57680 | Peptidase S41 family protein |
| K01236 | 898 | AT5G12080 | mechanosensitive channel of small conductance-like 10 |
| K01235 | 899 | AT1G51290 | F-box and associated interaction domains-containing protein |
| K01234 | 900 | AT5G35330 | methyl-CPG-binding domain protein 02 |
| K01233 | 901 | AT5G35330 | methyl-CPG-binding domain protein 02 |
| K01232 | 902 | AT4G29900 | autoinhibited Ca(2+)-ATPase 10 |
| K01231 | 903 | AT3G46920 | Protein kinase superfamily protein with octicosapeptide/Phox/Bem1p domain |
| K01230 | 904 | AT5G53390 | O-acyltransferase (WSD1-like) family protein |
| K01228/K01229 | 905 | AT2G26430 | arginine-rich cyclin 1 |
| K01227 | 907 | | |
| K01226 | 908 | AT5G18750 | DNAJ heat shock N-terminal domain-containing protein |
| K01225 | 909 | AT3G07610 | Transcription factor jumonji (jmjC) domain-containing protein |
| K01224 | 910 | AT5G65430 | general regulatory factor 8 |
| K01223 | 911 | AT5G55160 | small ubiquitin-like modifier 2 |
| K01222 | 912 | AT3G25040 | endoplasmic reticulum retention defective 2B |
| K01221 | 913 | AT4G24610 | |
| K01220 | 914 | AT3G49000 | RNA polymerase III subunit RPC82 family protein |
| K01219 | 915 | AT2G23360 | Plant protein of unknown function (DUF869) |
| K01218 | 916 | | |
| K01217 | 917 | AT3G07610 | Transcription factor jumonji (jmjC) domain-containing protein |
| K01216 | 918 | AT3G07610 | Transcription factor jumonji (jmjC) domain-containing protein |
| K01215 | 919 | | |
| K01214 | 920 | AT4G08620 | sulphate transporter 1;1 |
| K01213 | 921 | AT2G17700 | ACT-like protein tyrosine kinase family protein |

| | | | |
|---|---|---|---|
| K01212 | 922 | AT3G50120 | Plant protein of unknown function (DUF247) |
| K01211 | 923 | AT5G13870 | xyloglucan endotransglucosylase/hydrolase 5 |
| K01210 | 924 | AT3G43660 | Vacuolar iron transporter (VIT) family protein |
| K01209 | 925 | AT3G51510 | |
| K01208 | 926 | AT3G55510 | Noc2p family |
| K00944 | 927 | AT5G16370 | acyl activating enzyme 5 |
| K01154 | 928 | AT5G16370 | acyl activating enzyme 5 |
| K01155 | 929 | AT3G02700 | NC domain-containing protein-related |
| K01156 | 930 | AT5G63870 | serine/threonine phosphatase 7 |
| K01157 | 931 | AT1G21540 | AMP-dependent synthetase and ligase family protein |
| K01158 | 932 | AT5G37020 | auxin response factor 8 |
| K01159 | 933 | AT1G78690 | Phospholipid/glycerol acyltransferase family protein |
| K01160 | 934 | | |
| K01161 | 935 | AT5G65930 | kinesin-like calmodulin-binding protein (ZWICHEL) |
| K01162 | 936 | AT5G45890 | senescence-associated gene 12 |
| K01163 | 937 | AT5G50260 | Cysteine proteinases superfamily protein |
| K01164 | 938 | AT1G65730 | YELLOW STRIPE like 7 |
| K01165 | 939 | AT1G21970 | Histone superfamily protein |
| K01166 | 940 | AT2G38880 | nuclear factor Y, subunit B1 |
| K01167 | 941 | AT1G30300 | Metallo-hydrolase/oxidoreductase superfamily protein |
| K01168 | 942 | | |
| K01169 | 943 | AT5G45890 | senescence-associated gene 12 |
| K01170 | 944 | | |
| K01171 | 945 | AT3G05320 | O-fucosyltransferase family protein |
| K01172 | 946 | AT1G31260 | zinc transporter 10 precursor |
| K01173 | 947 | AT1G31260 | zinc transporter 10 precursor |
| K01174 | 948 | ATCG00680 | photosystem II reaction center protein B |
| K01175 | 949 | AT2G35035 | urease accessory protein D |
| K01176 | 950 | AT1G21970 | Histone superfamily protein |
| K01177 | 951 | AT1G24310 | |
| K01178 | 952 | AT5G16370 | acyl activating enzyme 5 |
| K01179 | 953 | AT2G31490 | |
| K01180 | 954 | AT1G65730 | YELLOW STRIPE like 7 |
| K01181 | 955 | | |
| K01182 | 956 | AT1G24350 | Acid phosphatase/vanadium-dependent haloperoxidase-related protein |
| K01183 | 957 | AT3G07610 | Transcription factor jumonji (jmjC) domain-containing protein |
| K01184 | 958 | AT1G78060 | Glycosyl hydrolase family protein |
| K01185 | 959 | AT5G36960 | |
| K01186 | 960 | ATCG00860 | Chloroplast Ycf2;ATPase, AAA type, core |
| K01187 | 961 | AT1G03700 | Uncharacterised protein family (UPF0497) |

| | | | |
|---|---|---|---|
| K01188 | 962 | AT3G02680 | nijmegen breakage syndrome 1 |
| K01189 | 963 | AT1G65690 | Late embryogenesis abundant (LEA) hydroxyproline-rich glycoprotein family |
| K01190 | 964 | AT1G10865 | |
| K01191 | 965 | AT3G48310 | cytochrome P450, family 71, subfamily A, polypeptide 22 |
| K01192 | 966 | AT5G47550 | Cystatin/monellin superfamily protein |
| K01193 | 967 | AT3G48310 | cytochrome P450, family 71, subfamily A, polypeptide 22 |
| K01194 | 968 | | |
| K01195 | 969 | AT4G16444 | |
| K01196 | 970 | AT1G21930 | |
| K01197 | 971 | | |
| K01198 | 972 | AT1G65700 | Small nuclear ribonucleoprotein family protein |
| K01199 | 973 | AT1G13320 | protein phosphatase 2A subunit A3 |
| K01200 | 974 | AT5G11380 | 1-deoxy-D-xylulose 5-phosphate synthase 3 |
| K01201 | 975 | AT3G02690 | nodulin MtN21 /EamA-like transporter family protein |
| K01202 | 976 | AT2G03200 | Eukaryotic aspartyl protease family protein |
| K01203 | 977 | AT1G65710 | |
| K01204 | 978 | AT1G65820 | microsomal glutathione s-transferase, putative |
| K01205 | 979 | AT3G07040 | NB-ARC domain-containing disease resistance protein |
| K01206 | 980 | AT5G61190 | putative endonuclease or glycosyl hydrolase with C2H2-type zinc finger domain |
| K01207 | 981 | AT5G04885 | Glycosyl hydrolase family protein |
| K00751 | 982 | ATCG00720 | photosynthetic electron transfer B |
| K00750 | 983 | AT1G29930 | chlorophyll A/B binding protein 1 |
| K00749 | 984 | AT1G73370 | sucrose synthase 6 |
| K00748 | 985 | AT5G57500 | Galactosyltransferase family protein |
| K00747 | 986 | AT1G65900 | |
| K00746 | 987 | AT3G48030 | hypoxia-responsive family protein / zinc finger (C3HC4-type RING finger) family protein |
| K00745 | 988 | AT1G65910 | NAC domain containing protein 28 |
| K00744 | 989 | AT1G78380 | glutathione S-transferase TAU 19 |
| K00743 | 990 | AT5G17290 | autophagy protein Apg5 family |
| K00742 | 991 | AT5G37260 | Homeodomain-like superfamily protein |
| K00741 | 992 | AT3G51520 | diacylglycerol acyltransferase family |
| K00740 | 993 | AT3G03250 | UDP-GLUCOSE PYROPHOSPHORYLASE 1 |
| K00739 | 994 | AT1G73360 | homeodomain GLABROUS 11 |
| K00738 | 995 | AT1G73360 | homeodomain GLABROUS 11 |
| K00737 | 996 | AT1G28690 | Tetratricopeptide repeat (TPR)-like superfamily protein |
| K00752 | 997 | AT3G17480 | F-box and associated interaction domains-containing protein |
| K00753 | 998 | | |
| K00754 | 999 | AT1G13690 | ATPase E1 |

| | | | |
|---|---|---|---|
| K00755 | 1000 | | |
| K00756 | 1001 | | |
| K00757 | 1002 | | |
| K00758 | 1003 | AT3G19760 | eukaryotic initiation factor 4A-III |
| K00759 | 1004 | AT5G63520 | |
| K00970 | 1005 | AT1G21270 | wall-associated kinase 2 |
| K00971 | 1006 | AT2G32090 | Lactoylglutathione lyase / glyoxalase I family protein |
| K00972 | 1007 | AT1G21240 | wall associated kinase 3 |
| K00973 | 1008 | AT1G21230 | wall associated kinase 5 |
| K00974 | 1009 | ATCG01310 | ribosomal protein L2 |
| K00975 | 1010 | AT5G17240 | SET domain group 40 |
| K00976 | 1011 | AT5G17230 | PHYTOENE SYNTHASE |
| K00977 | 1012 | AT5G36110 | cytochrome P450, family 716, subfamily A, polypeptide 1 |
| K00978 | 1013 | AT5G17220 | glutathione S-transferase phi 12 |
| K00979 | 1014 | AT1G65890 | acyl activating enzyme 12 |
| K00980 | 1015 | AT4G19640 | Ras-related small GTP-binding family protein |
| K00981 | 1016 | AT1G65840 | polyamine oxidase 4 |
| K00982 | 1017 | AT3G02720 | Class I glutamine amidotransferase-like superfamily protein |
| K00983 | 1018 | AT3G02710 | ARM repeat superfamily protein |
| K01079 | 1019 | AT1G73360 | homeodomain GLABROUS 11 |
| K01080 | 1020 | | |
| K01081 | 1021 | | |
| K01082 | 1022 | | |
| K01083 | 1023 | | |
| K01084 | 1024 | | |
| K01085 | 1025 | AT1G65950 | Protein kinase superfamily protein |
| K01086 | 1026 | AT1G65930 | cytosolic NADP+-dependent isocitrate dehydrogenase |
| K01087 | 1027 | AT1G73360 | homeodomain GLABROUS 11 |
| K01088 | 1028 | AT1G73360 | homeodomain GLABROUS 11 |
| K01089 | 1029 | AT3G03480 | acetyl CoA:(Z)-3-hexen-1-ol acetyltransferase |
| K01090 | 1030 | | |
| K01091 | 1031 | | |
| K01092 | 1032 | AT1G73360 | homeodomain GLABROUS 11 |
| K01093 | 1033 | AT3G03480 | acetyl CoA:(Z)-3-hexen-1-ol acetyltransferase |
| K01094 | 1034 | AT5G17540 | HXXXD-type acyl-transferase family protein |
| K01095 | 1035 | AT3G03480 | acetyl CoA:(Z)-3-hexen-1-ol acetyltransferase |
| K01096 | 1036 | AT3G03270 | Adenine nucleotide alpha hydrolases-like superfamily protein |
| K01097 | 1037 | AT3G03270 | Adenine nucleotide alpha hydrolases-like superfamily protein |
| K01098 | 1038 | AT1G51790 | Leucine-rich repeat protein kinase family protein |
| K01099 | 1039 | | |

| | | | |
|---|---|---|---|
| K01100 | 1040 | AT3G03480 | acetyl CoA:(Z)-3-hexen-1-ol acetyltransferase |
| K01101 | 1041 | | |
| K01102 | 1042 | AT5G17540 | HXXXD-type acyl-transferase family protein |
| K01103 | 1043 | AT5G17540 | HXXXD-type acyl-transferase family protein |
| K01104 | 1044 | AT3G03270 | Adenine nucleotide alpha hydrolases-like superfamily protein |
| K01105 | 1045 | AT5G17540 | HXXXD-type acyl-transferase family protein |
| K01115 | 1046 | AT5G17540 | HXXXD-type acyl-transferase family protein |
| K01114 | 1047 | AT4G05230 | Ubiquitin-like superfamily protein |
| K01113 | 1048 | AT5G37370 | PRP38 family protein |
| K01112 | 1049 | AT5G37478 | TPX2 (targeting protein for Xklp2) protein family |
| K01111 | 1050 | AT1G66080 | |
| K01110 | 1051 | AT5G16550 | |
| K01109 | 1052 | AT1G35210 | |
| K01108 | 1053 | | |
| K01107 | 1054 | AT5G16560 | Homeodomain-like superfamily protein |
| K01106 | 1055 | AT1G70520 | cysteine-rich RLK (RECEPTOR-like protein kinase) 2 |
| K00943 | 1056 | AT3G23220 | Integrase-type DNA-binding superfamily protein |
| K00942 | 1057 | AT3G23240 | ethylene response factor 1 |
| K00941 | 1058 | AT3G23240 | ethylene response factor 1 |
| K00940 | 1059 | AT3G23240 | ethylene response factor 1 |
| K00939 | 1060 | AT3G23240 | ethylene response factor 1 |
| K00938 | 1061 | AT4G01240 | S-adenosyl-L-methionine-dependent methyltransferases superfamily protein |
| K00937 | 1062 | AT3G02850 | STELAR K+ outward rectifier |
| K00936 | 1063 | AT3G02860 | zinc ion binding |
| K00935 | 1064 | AT3G07530 | |
| K00934 | 1065 | AT5G37530 | NAD(P)-binding Rossmann-fold superfamily protein |
| K00933 | 1066 | AT3G52640 | Zn-dependent exopeptidases superfamily protein |
| K00932 | 1067 | AT3G18670 | Ankyrin repeat family protein |
| K00931 | 1068 | AT1G21970 | Histone superfamily protein |
| K00930 | 1069 | AT2G24840 | AGAMOUS-like 61 |
| K00929 | 1070 | ATCG00720 | photosynthetic electron transfer B |
| K00928 | 1071 | AT3G56830 | Protein of unknown function (DUF565) |
| K00927 | 1072 | AT5G18525 | protein serine/threonine kinases;protein tyrosine kinases;ATP binding;protein kinases |
| K00926 | 1073 | AT5G37510 | NADH-ubiquinone dehydrogenase, mitochondrial, putative |
| K00925 | 1074 | AT5G37540 | Eukaryotic aspartyl protease family protein |
| K00924 | 1075 | AT1G09680 | Pentatricopeptide repeat (PPR) superfamily protein |
| K00923 | 1076 | AT5G66460 | Glycosyl hydrolase superfamily protein |
| K00922 | 1077 | AT5G66460 | Glycosyl hydrolase superfamily protein |
| K00921 | 1078 | | |

| | | | |
|---|---|---|---|
| K00920 | 1079 | AT5G37600 | glutamine synthase clone R1 |
| K00919 | 1080 | | |
| K00918 | 1081 | AT5G37630 | ARM repeat superfamily protein |
| K00917 | 1082 | AT2G01670 | nudix hydrolase homolog 17 |
| K00916 | 1083 | AT5G01750 | Protein of unknown function (DUF567) |
| K00915 | 1084 | AT3G18150 | RNI-like superfamily protein |
| K00914 | 1085 | AT5G37670 | HSP20-like chaperones superfamily protein |
| K00913 | 1086 | AT5G17190 | |
| K00912 | 1087 | | |
| K00911 | 1088 | AT1G61150 | LisH and RanBPM domains containing protein |
| K00910 | 1089 | AT1G48480 | receptor-like kinase 1 |
| K00909 | 1090 | AT1G48480 | receptor-like kinase 1 |
| K00908 | 1091 | | |
| K00907 | 1092 | AT5G66460 | Glycosyl hydrolase superfamily protein |
| K00906 | 1093 | AT5G66460 | Glycosyl hydrolase superfamily protein |
| K00905 | 1094 | AT5G16600 | myb domain protein 43 |
| K00904 | 1095 | AT5G16610 | |
| K00903 | 1096 | AT1G66250 | O-Glycosyl hydrolases family 17 protein |
| K00902 | 1097 | AT5G37710 | alpha/beta-Hydrolases superfamily protein |
| K00901 | 1098 | AT1G59720 | Tetratricopeptide repeat (TPR)-like superfamily protein |
| K00900 | 1099 | AT1G14900 | high mobility group A |
| K00899 | 1100 | AT5G16720 | Protein of unknown function, DUF593 |
| K00898 | 1101 | AT5G46800 | Mitochondrial substrate carrier family protein |
| K00897 | 1102 | AT5G28040 | DNA-binding storekeeper protein-related transcriptional regulator |
| K00896 | 1103 | AT5G16710 | dehydroascorbate reductase 1 |
| K00895 | 1104 | AT2G18620 | Terpenoid synthases superfamily protein |
| K00894 | 1105 | AT1G14920 | GRAS family transcription factor family protein |
| K00893 | 1106 | AT1G09910 | Rhamnogalacturonate lyase family protein |
| K00892 | 1107 | AT1G19320 | Pathogenesis-related thaumatin superfamily protein |
| K00891 | 1108 | AT3G06240 | F-box family protein |
| K00890 | 1109 | | |
| K00889 | 1110 | AT4G39330 | cinnamyl alcohol dehydrogenase 9 |
| K00888 | 1111 | AT4G37990 | elicitor-activated gene 3-2 |
| K00888 | 1111 | AT4G39330 | cinnamyl alcohol dehydrogenase 9 |
| K00887 | 1112 | AT2G02820 | myb domain protein 88 |
| K00886 | 1113 | | |
| K00809 | 1114 | AT3G13230 | RNA-binding KH domain-containing protein |
| K00808 | 1115 | AT3G13230 | RNA-binding KH domain-containing protein |
| K00807 | 1116 | AT5G65530 | Protein kinase superfamily protein |
| K00806 | 1117 | AT5G65520 | Tetratricopeptide repeat (TPR)-like superfamily protein |

| | | | |
|---|---|---|---|
| K00805 | 1118 | AT5G57390 | AINTEGUMENTA-like 5 |
| K00804 | 1119 | AT4G38050 | Xanthine/uracil permease family protein |
| K00803 | 1120 | AT5G65495 | |
| K00802 | 1121 | AT5G65490 | |
| K00801 | 1122 | AT2G22570 | nicotinamidase 1 |
| K00800 | 1123 | AT2G22560 | Kinase interacting (KIP1-like) family protein |
| K00799 | 1124 | AT4G38060 | |
| K00798 | 1125 | AT4G38070 | basic helix-loop-helix (bHLH) DNA-binding superfamily protein |
| K00797 | 1126 | AT5G10490 | MSCS-like 2 |
| K00796 | 1127 | AT4G24530 | O-fucosyltransferase family protein |
| K00795 | 1128 | | |
| K00794 | 1129 | AT1G09820 | Pentatricopeptide repeat (PPR-like) superfamily protein |
| K00793 | 1130 | AT5G10470 | kinesin like protein for actin based chloroplast movement 1 |
| K00792 | 1131 | AT2G22530 | Alkaline-phosphatase-like family protein |
| K00791 | 1132 | AT1G62970 | Chaperone DnaJ-domain superfamily protein |
| K00790 | 1133 | | |
| K00789 | 1134 | | |
| K00788 | 1135 | | |
| K00787 | 1136 | | |
| K00786 | 1137 | AT4G38090 | Ribosomal protein S5 domain 2-like superfamily protein |
| K00785 | 1138 | AT5G65450 | ubiquitin-specific protease 17 |
| K00784 | 1139 | AT2G22500 | uncoupling protein 5 |
| K00783 | 1140 | | |
| K00782 | 1141 | | |
| K00781 | 1142 | AT5G38190 | |
| K00780 | 1143 | AT1G68960 | Protein of unknown function (DUF295) |
| K00779 | 1144 | AT4G24610 | |
| K00778 | 1145 | AT2G39445 | Phosphatidylinositol N-acetylglucosaminyltransferase, GPI19/PIG-P subunit |
| K00777 | 1146 | AT1G80770 | P-loop containing nucleoside triphosphate hydrolases superfamily protein |
| K00776 | 1147 | AT5G65430 | general regulatory factor 8 |
| K00775 | 1148 | AT5G65420 | CYCLIN D4;1 |
| K00774 | 1149 | AT2G22480 | phosphofructokinase 5 |
| K00773 | 1150 | | |
| K00772 | 1151 | AT1G26290 | |
| K00771 | 1152 | AT2G23360 | Plant protein of unknown function (DUF869) |
| K00770 | 1153 | AT3G49000 | RNA polymerase III subunit RPC82 family protein |
| K00769 | 1154 | AT1G45688 | |
| K00768 | 1155 | AT5G67080 | mitogen-activated protein kinase kinase kinase 19 |
| K00767 | 1156 | AT5G67030 | zeaxanthin epoxidase (ZEP) (ABA1) |

| | | | |
|---|---|---|---|
| K00766 | 1157 | AT5G67020 | |
| K00765 | 1158 | AT1G18210 | Calcium-binding EF-hand family protein |
| K00764 | 1159 | AT3G19200 | |
| K00763 | 1160 | AT3G19184 | AP2/B3-like transcriptional factor family protein |
| K00762 | 1161 | AT3G19150 | KIP-related protein 6 |
| K00761 | 1162 | AT3G05090 | Transducin/WD40 repeat-like superfamily protein |
| K00760 | 1163 | | |
| K00816 | 1164 | AT3G50620 | P-loop containing nucleoside triphosphate hydrolases superfamily protein |
| K00817 | 1165 | AT3G50530 | CDPK-related kinase |
| K00818 | 1166 | AT2G26560 | phospholipase A 2A |
| K00819 | 1167 | | |
| K00820 | 1168 | | |
| K00821 | 1169 | | |
| K00822 | 1170 | | |
| K00823 | 1171 | | |
| K00824 | 1172 | | |
| K00825 | 1173 | AT5G66850 | mitogen-activated protein kinase kinase kinase 5 |
| K00826 | 1174 | AT5G66860 | Ribosomal protein L25/Gln-tRNA synthetase, anti-codon-binding domain |
| K00827 | 1175 | AT2G23660 | LOB domain-containing protein 10 |
| K00828 | 1176 | AT4G37180 | Homeodomain-like superfamily protein |
| K00829 | 1177 | AT5G66880 | sucrose nonfermenting 1(SNF1)-related protein kinase 2.3 |
| K00830 | 1178 | AT4G37170 | Pentatricopeptide repeat (PPR) superfamily protein |
| K00831 | 1179 | AT5G66920 | SKU5 similar 17 |
| K00832 | 1180 | AT3G50440 | methyl esterase 10 |
| K00833 | 1181 | AT3G50410 | OBF binding protein 1 |
| K00834 | 1182 | AT2G23540 | GDSL-like Lipase/Acylhydrolase superfamily protein |
| K00835 | 1183 | AT4G37100 | Pyridoxal phosphate (PLP)-dependent transferases superfamily protein |
| K00836 | 1184 | AT2G23470 | Protein of unknown function, DUF647 |
| K00837 | 1185 | AT2G26490 | Transducin/WD40 repeat-like superfamily protein |
| K00838 | 1186 | AT4G37090 | |
| K00839 | 1187 | AT2G23450 | Protein kinase superfamily protein |
| K00840 | 1188 | AT4G37200 | Thioredoxin superfamily protein |
| K00841 | 1189 | AT1G10747 | Maternally expressed gene (MEG) family protein |
| K00842 | 1190 | AT4G37210 | Tetratricopeptide repeat (TPR)-like superfamily protein |
| K00843 | 1191 | | |
| K00844 | 1192 | AT3G50810 | Uncharacterised protein family (UPF0497) |
| K00845 | 1193 | AT2G29760 | Tetratricopeptide repeat (TPR)-like superfamily protein |
| K00846 | 1194 | | |
| K00847 | 1195 | AT2G23755 | |

| | | | |
|---|---|---|---|
| K00848 | 1196 | | |
| K00849 | 1197 | AT3G50780 | |
| K00850 | 1198 | AT4G36860 | LIM domain-containing protein |
| K00851 | 1199 | AT4G36860 | LIM domain-containing protein |
| K00852 | 1200 | AT5G66631 | Tetratricopeptide repeat (TPR)-like superfamily protein |
| K00853 | 1201 | AT2G33580 | Protein kinase superfamily protein |
| K00854 | 1202 | AT1G19690 | NAD(P)-binding Rossmann-fold superfamily protein |
| K00855 | 1203 | AT2G23790 | Protein of unknown function (DUF607) |
| K00856 | 1204 | AT4G34320 | Protein of unknown function (DUF677) |
| K00857 | 1205 | AT1G13245 | ROTUNDIFOLIA like 17 |
| K00858 | 1206 | AT1G67260 | TCP family transcription factor |
| K00859 | 1207 | AT5G38660 | acclimation of photosynthesis to environment |
| K00860 | 1208 | AT5G38640 | NagB/RpiA/CoA transferase-like superfamily protein |
| K00861 | 1209 | AT5G38630 | cytochrome B561-1 |
| K00862 | 1210 | | |
| K00863 | 1211 | AT3G02420 | |
| K00864 | 1212 | AT5G38560 | Protein kinase superfamily protein |
| K00865 | 1213 | AT5G15900 | TRICHOME BIREFRINGENCE-LIKE 19 |
| K00866 | 1214 | AT5G15890 | TRICHOME BIREFRINGENCE-LIKE 21 |
| K00867 | 1215 | AT1G14130 | 2-oxoglutarate (2OG) and Fe(II)-dependent oxygenase superfamily protein |
| K00868 | 1216 | AT1G14130 | 2-oxoglutarate (2OG) and Fe(II)-dependent oxygenase superfamily protein |
| K00869 | 1217 | AT1G14130 | 2-oxoglutarate (2OG) and Fe(II)-dependent oxygenase superfamily protein |
| K00870 | 1218 | AT3G30841 | Cofactor-independent phosphoglycerate mutase |
| K00871 | 1219 | AT5G38710 | Methylenetetrahydrofolate reductase family protein |
| K00872 | 1220 | AT5G38720 | |
| K00873 | 1221 | AT5G38760 | Late embryogenesis abundant protein (LEA) family protein |
| K00874 | 1222 | AT2G24762 | glutamine dumper 4 |
| K00875 | 1223 | AT3G02350 | galacturonosyltransferase 9 |
| K00876 | 1224 | AT4G18430 | RAB GTPase homolog A1E |
| K00877 | 1225 | AT3G30530 | basic leucine-zipper 42 |
| K00878 | 1226 | AT5G15680 | ARM repeat superfamily protein |
| K00879 | 1227 | AT5G49550 | |
| K00880 | 1228 | AT5G15700 | DNA/RNA polymerases superfamily protein |
| K00881 | 1229 | AT2G33730 | P-loop containing nucleoside triphosphate hydrolases superfamily protein |
| K00882 | 1230 | AT5G15750 | Alpha-L RNA-binding motif/Ribosomal protein S4 family protein |
| K00883 | 1231 | AT5G15780 | Pollen Ole e 1 allergen and extensin family protein |
| K00884 | 1232 | AT3G45080 | P-loop containing nucleoside triphosphate hydrolases superfamily protein |

| | |
|---|---|
| K00885 | 1233 |

Table S5. Inter- and intra-population nucleotide diversity levels comparing CenH3A[a] to other regions on LG14.

| Comparison[b] | Mean $\pi$[c] | Mean of permuted means[d] | Range of permuted means | Standard deviation of permuted means | *P*[e] |
|---|---|---|---|---|---|
| IM-IM | 0.00232 | 0.00796 | (0.00109 - 0.01920) | 0.00014 | **0.016** |
| IM-AHQT | 0.00413 | 0.00909 | (0.00172 - 0.02229) | 0.00017 | 0.086 |
| IM-DUN | 0.00488 | 0.00708 | (0.00088 - 0.01407) | 0.00011 | 0.204 |
| IM-LMC24 | 0.00630 | 0.00899 | (0.00070 - 0.01945) | 0.00014 | 0.224 |
| IM-MAR3 | 0.00461 | 0.00920 | (0.00098 - 0.02331) | 0.00018 | 0.116 |

[a]Diversity values computed for 8-gene block surrounding CenH3A

[b] 34 IM lines and a line from a distant population (Table S1)

[c] Nei's diversity per gene per site (Nei 1979)

[d]500 permutations were performed by averaging π per site per gene from blocks of 8 consecutive genes along LG14

[e]P values generated by comparing estimated means to permuted means

Table S6. Seven distinct, long-range haplotypes with more than one individual define the CenH3A region of LG14. Start and end coordinates on LG14, size in base pairs, IM individuals with the haplotype, and the number of individuals are given for haplotypes 1-7, as well as the identities of four singleton lines with unique haplotypes.

| Haplotype | Start position | End position | Size (bp) | Individuals | N |
|---|---|---|---|---|---|
| 1 | 13836336 | 13581975 | 254361 | IM1054, IM239, IM275, IM170, IM767, IM835, IM742 | 7 |
| 2 | 13105111 | 13725184 | 620073 | IM667, IM664, IM1192 | 3 |
| 3 | 13770309 | 13551509 | 218800 | IM549, IM359 | 2 |
| 4 | 13570360 | 13738836 | 168476 | IM624, IM62, IM179, IM693, IM777, IM479, IM709 | 7 |
| 5 | 13581975 | 13720468 | 138493 | IM109, IM922, IM238, IM115, IM909, IM657 | 6 |
| 6 | 13555923 | 13720169 | 164246 | IM1145, IM785, IM116 | 3 |
| 7 | 13506401 | 13902746 | 396345 | IM412, IM138 | 2 |
| singletons | N/A | N/A | N/A | IM266, IM502, IM1152, IM106 | 1 each |

Table S7. Marker names, genes, primer sequences, and product sizes for genetic markers used in this study.

| Marker | Gene | Forward Primer | Reverse Primer | Ref allele size( bp) |
|---|---|---|---|---|
| lb5a | Migut.K00858 | CGGAGAATATATCGTGGTGG | ACTGCACCTCTCAATCTTGG | 276 |
| mK1229/J575 | Migut.K01229/ Migut.J00575 | TGTGGATCTAAAGGGAGATTTGA | TCATTTGCAAGATTCCATGC | 173 /180 |
| mCenH3A | Migut.N01557 | AAGAAATCCTCCGGTGAGAA | AACATGGTGTAGCAGTTGTGC | 274 |
